# Gene expression dynamics through time reveal low pleiotropy genes underlying local adaptation in silver birch (*Betula pendula*)

**DOI:** 10.64898/2026.10.16.759317

**Authors:** Jenni Sirén, Jaana Vuosku, Sonja T. Kujala, Katri Kärkkäinen, Ysaline Perraud, Jenni Punttila, Jukka-Pekka Verta, Tanja Pyhäjärvi

**Author notes:** Author for correspondence: Jenni Sirén.

## Abstract

Understanding the molecular mechanisms underlying local adaptation in boreal trees is important for assessing their adaptive potential under climate change. The role of pleiotropy in polygenic local adaptation remains poorly understood. Here we studied growth and temporal transcriptomic responses to photoperiod in three photoperiod treatments in three populations of *Betula pendula* within a two-week time period to identify genes and co-expression networks underlying local adaptation and to estimate the level of pleiotropy in population specific gene expression. We used RNA sequencing to characterize the temporal transcriptomic responses which were analyzed with variance partitioning, differential gene expression and weighted co-expression network analyses. Photoperiod treatment and sampling time were the main drivers of gene expression variation both at the co-expression module and single gene level. The majority of circadian clock and flowering time genes exhibited striking similarity across populations. Nevertheless, we detected small but significant population effects in the temporal gene expression patterns which were concentrated on the periphery of the co-expression networks consistent with low pleiotropy effects. Our findings suggest that local adaptation to growing season length among populations exhibiting very low genetic differentiation along a latitudinal cline in *B. pendula* is likely polygenic and potentially governed through low pleiotropy genes.

## Introduction

Plants as sessile organisms depend on their ability to adapt to the surrounding environments for survival. Populations in species whose range overlaps heterogeneous environments are typically adapted to the local conditions (Kawecki & Ebert, 2004; Savolainen *et al*., 2013) and in species with large continuous ranges this adaptation is often clinal in relation to latitude, longitude or altitude (Endler, 1977; Savolainen *et al*., 2007; Aitken *et al*., 2008; Alberto *et al*., 2013a). For trees in boreal forests, photoperiod is the most reliable environmental cue to time their phenology, such as growth cessation and reproduction (Garner & Allard, 1920; Heide, 1974; Petterle *et al*., 2013) and differences in phenology among populations, especially at the end of growing season as a response to photoperiod are documented in several widely distributed, ecologically significant trees such as *Pinus sylvestris* (Savolainen et al., 2004), *Quercus petraea* (Vitasse et al., 2009), and *Betula pendula* (Li et al., 2003; Viherä-Aarnio et al., 2005).

Phenological responses have been extensively studied in plants, resulting in a substantial body of knowledge on the genes and pathways mediating responses to photoperiod under both natural and controlled light conditions (McClung, 2001; Böhlenius *et al*., 2006; Slotte *et al*., 2007; Ibáñez *et al*., 2010; de Montaigu *et al*., 2015; Lázaro-Gimeno *et al*., 2024; Mariën *et al*., 2025). In particular, circadian clock and flowering-time genes have been shown to be targets of natural selection in e.g., *Populus* species (Hall *et al*., 2011; Keller *et al*., 2012; Evans *et al*., 2014) and *Arabidopsis lyrata* (Mattila *et al*., 2016). Population-level differences in gene expression have been reported in e.g., conifer species in Northern Europe indicating local adaptation potentially proceeding through gene regulation. In northern Finnish populations of *Pinus sylvestris* the expression of *FLOWERING LOCUS T/TERMINAL FLOWER 1*-like (*FTL2*) gene was higher under long (16 h) nights compared to southern Finnish and Polish populations (Avia et al., 2014). In *Picea abies*, *FTL2* had a higher expression level under 8 h nights in northern Swedish populations compared to southern Swedish populations (Chen *et al*., 2012). However, since the molecular mechanisms of photoperiodic responses in plants have been of great interest and individual loci were characterized relatively early (Gendron & Staiger, 2023), they have widely been *a priori* candidates of adaptation studies potentially leaving other locally adaptive loci and mechanisms understudied.

Gene expression differences link environmental cues to adaptive genetic variation among populations (Whitehead & Crawford, 2006; Fraser, 2013; Gould *et al*., 2018; Blanc *et al*., 2021; Tisinai *et al*., 2026). The expected relationship is not always one-to-one, where adaptive genetic differences in a given gene or in its regulatory region would impact mainly that gene’s expression and vice versa. Rather, even a single genetic change has the potential to affect the whole gene regulatory network. Based on this, cost of complexity theory (Orr, 2000; Wagner & Zhang, 2011) postulates that genes that influence a large number of other genes, which are typically central hub genes in gene co-expression networks, would have a constraint for adaptation compared to low pleiotropy genes which usually are peripheral in the networks.

In contrast to this prediction, Whiting *et al*., 2024 found that across many plant species, repeated local adaptation was mediated mainly through high pleiotropy genes. In addition, both Hämälä *et al*., 2020 and Okinedo and Moyers, 2026 found that high pleiotropy genes are governing local adaptation in *Ambrosia artemisiifolia* and *Helianthus argophyllus*. However, the study on repeated adaptation may not reflect overall patterns of local adaptation at the species level. In the latter two studies, the optima of different populations were far away from each other, which also does not represent the situation in clinal local adaptation. In the case of forest trees there are only few examples of single genes showing allele frequency differences consistent with strong clinal selection (but see Ingvarsson *et al*., 2006), because adaptation is likely often polygenic (Barghi *et al*., 2020) and individual adaptive loci are challenging to pinpoint based on genetic data alone (Le Corre & Kremer, 2012; Yeaman, 2015; Höllinger *et al*., 2019). This leaves a knowledge-gap in our understanding of the level of pleiotropy in cases of polygenic adaptation, such as seen in forest trees.

Silver birch (*B. pendula*) is an ecologically and economically important forest tree species in Northern Europe. It is outcrossing, wind-pollinated, and has a large, continuous distribution resulting in low genetic differentiation among populations (FST among typical Nordic populations < 0.01 (Milesi et al., 2024; Herrera-Egoavil et al., 2026)). It exhibits clinal local adaptation to growing season length at the phenotypic level across latitudes. Timing of growth cessation depends on the latitude of seed origin in first year seedlings (Viherä-Aarnio *et al*., 2005) and the critical photoperiod and photoperiodic sensitivity for growth cessation vary with latitudinal origin of the population (Li *et al*., 2003). However, the molecular mechanisms underlying these adaptations are not well understood. Based on whole genome resequencing, (Salojärvi *et al*., 2017) found signatures of selective sweeps and a group of six *Arabidopsis thaliana* homologs (*SWEETIE*, *KAKTUS* (*KAK*), *ARABIDOPSIS RESPONSE REGULATOR 1* (*ARR1*), *MED5A*, *PHYTOCHROME C* (*PHYC*), and *FAR1-RELATED SEQUENCE 10* (*FRS10*)) associated with environmental factors in a range-wide sample of *B. pendula*. Recently, using exome capture and high-density sampling across a latitudinal gradient in Sweden, Herrera-Egoavil *et al*., 2026 found that the genetic signal for local adaptation was mostly concentrated in an inversion in chromosome 1. However, the role of gene expression and its regulation through time in relation to the environmental cues, and their contribution to local adaptation have not yet been investigated at the transcriptome-wide level in *B. pendula*.

Here, we analysed leaf transcriptomes of *B. pendula* seedlings originating from different populations subjected to three different photoperiod treatments. Seedling growth and transcriptomes were monitored across time to allow detection of potential population-specific responses to photoperiod. A combination of variance partitioning, co-expression network analysis and linear mixed-effect models allowed evaluation of the relative impact of local adaptation to gene expression patterns, as well as testing of cost-of-complexity theory in a clinal setting where population optima are close to each other and the populations are connected by extensive gene flow.

## Materials and Methods

### Controlled condition growth chamber experiments

We used open-pollinated *Betula pendula* Roth seeds (half-sib families) from eight trees from each of the four study populations (Oravainen (ORA, 63°18′N, 22°22′E), Polvijärvi (POL, 62°51′N, 29°22′E), Punkaharju (PUN, 61°45′N, 29°23′ E) and Vallerstad (VAL, 58°24′N, 15°06′E), Fig. 1a). Seeds were germinated for two weeks on moist tissue paper in Petri dishes under an 18 h : 6 h light : dark photoperiod at 22 °C. Seedlings were transplanted into Plantek PL64F seedling trays, with two seeds from each mother tree per tray (64 seedlings per tray). Ten trays were placed in each of three growth chambers (30 trays, 1920 seedlings in total). Seedling positions within trays were randomized, and trays were rotated weekly within chambers to reduce positional effects.

**Fig. 1.**
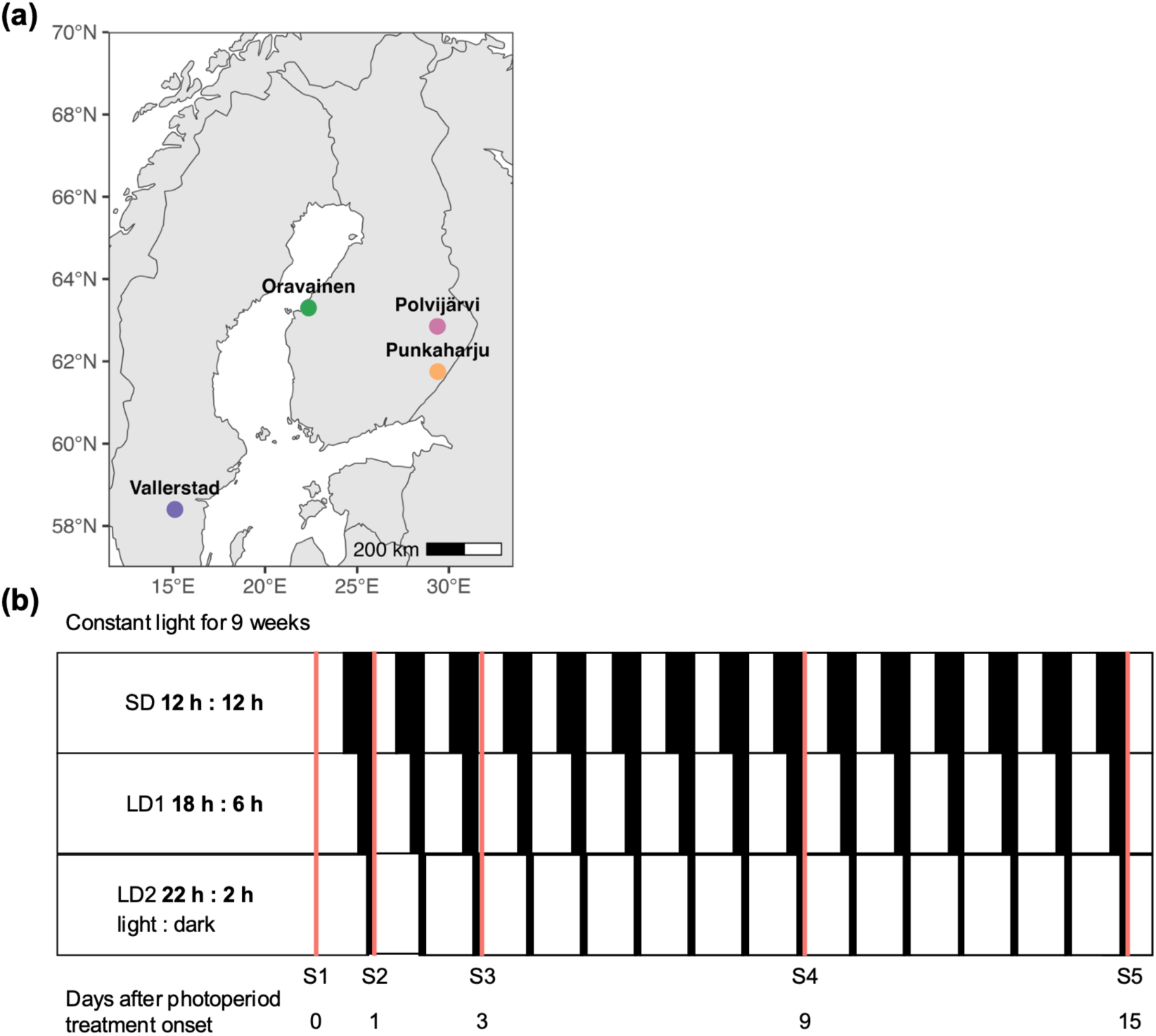
(a) Geographic locations of the *Betula pendula* populations used in this study. (b) Experimental design. Each row represents a growth chamber, black rectangles represent the night length, and vertical red lines represent the sampling time points.

Seedlings were grown in growth chambers at 20°C in constant light (CL) for nine weeks after which photoperiod treatments were applied at the chamber level to trigger population level differences (Fig. 1b). The three photoperiod treatments were 12 h : 12 h (SD), 18 h : 6 h (LD1) and 22 h : 2 h (LD2) (light : dark). The experiment was run in growth chambers with fixed light intensity (200 µmol m^−2^ s^−1^) and spectrum.

The first sampling (S1) was conducted under constant light prior to the onset of photoperiod treatments. After photoperiod initiation, the samplings were conducted immediately after the first dark period (S2) and then after 3 (S3), 9 (S4), and 15 (S5) days. Sampling was conducted within a two-hour period following the dark period. At each time point, two randomly selected trays per chamber were sampled, corresponding to four half-sib seedlings per family from each photoperiod treatment.

Sampling was destructive; thus, it was not possible to sample the same seedling more than once. For each sampled seedling, 2-3 of the uppermost leaves were collected into 2 ml Eppendorf tubes, flash-frozen in liquid nitrogen and stored at −80°C. Seedling height was measured first for all individuals nine days before initiating the photoperiod treatment and the second measurement was taken from the selected seedlings immediately prior to sampling.

### Growth data

We used linear mixed-effects models to analyse seedling growth. Growth was measured as absolute growth rate (AGR; Δgrowth/Δdays, mm/day). First, we fitted a full interaction model including sampling time, treatment, population and all possible interactions as fixed effects and mother tree as a cluster variable to account for genetic relatedness within populations. To reduce collinearity between main effects and interaction terms and to improve parameter interpretability, the continuous covariate (sampling time) was mean centred prior to model fitting. The models were fitted using the lmer() function from the lme4 package (Bates *et al*., 2015) in R. Model simplification was performed using backward stepwise selection with the step() function from the lmerTest package (Kuznetsova *et al*., 2017). Non-significant fixed effects and interactions were sequentially removed based on F-tests with Satterthwaite’s approximation for denominator degrees of freedom, resulting in the final model. In addition, the full and final models were compared using AIC and BIC (Table S1). The final model was specified as:

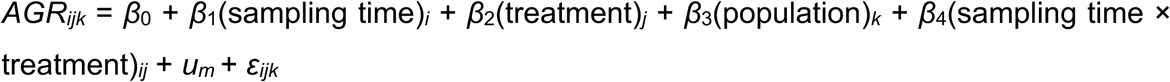

where *β* represents the fixed effect coefficients, *um* is the random intercept for the *m*-th mother tree (*um* ∼ *N*(0, *σ*^2^*mother*)), and *εijk* is the residual error.

### RNA extraction and sequencing

A subset of 214 samples consisting of three populations (Oravainen, Punkaharju and Vallerstad, Fig. 1a), 5-6 families from each population, all three photoperiod treatments and four sampling points (S1-S4) were used for RNA-seq. The frozen leaf samples were homogenized using TissueLyzer II (Qiagen) with 25 Hz shaking in 30 second intervals. Samples were kept on dry ice, dipped in liquid nitrogen before homogenization and transferred back onto dry ice immediately after homogenization. Total RNA was extracted using the Sigma Spectrum Plant Total RNA kit (cat.nro STRN250) Protocol B with the On Column DNaseI Digestion module (cat.nro DNASE70). Deviating from the standard protocol, 2.5 % (v/w) of PVP40 was added to the lysis buffer. For highly viscous samples, the lysis buffer volume was increased to 700 µl and the binding buffer volume to 350 µl. The concentration and purity of the RNA was assessed with Nanodrop 2000 (ThermoFisher) and Qubit 4.0 (ThermoFisher) using the RNA HS Assay Kit (cat.nro Q32855). Samples were sent to Biomarker Technologies (BMKGene) for library preparation and mRNA sequencing. Paired-end sequencing was conducted with Illumina NovaSeq X platform.

### Read quality, mapping to reference genome and counting reads mapped to genes

The initial quality control and trimming of adapters and low-quality reads was performed using fastp (version 0.21.0) (Chen, 2025) with options --detect_adapter_for_pe -- length_required 50 --cut_mean_quality 20 --cut_front --cut_tail --trim_front1=2 -- trim_front2=2. The reads were mapped to the *B. pendula* reference genome (Ord *et al*., unpublished) using STAR (version 2.7.11b) (Dobin *et al*., 2013) with default parameters. Additionally, the reads were mapped to the *B. pubescens* (Ord *et al*., unpublished) reference genome as a confirmation of species identity.

The number of reads mapped to genes was counted with featureCounts (version 2.0.8) (Liao *et al*., 2014), using -p --countReadPairs -M --fraction -s 2 parameters. The produced expression matrix was used in the subsequent analyses. MultiQC (version 1.25.1) (Ewels *et al*., 2016) was used to compare all reports.

Low-count genes were first filtered out by retaining only genes with a minimum count of 10 in at least six samples (n = 24662 genes). Variance-stabilizing transformation (VST) was then applied to the expression count matrix using the vst-function from the DESeq2 R package (Love *et al*., 2014) in R and the normalized count matrix was used in all subsequent analyses unless otherwise indicated. The sample relationships were assessed by hierarchical clustering of the expression matrix using Euclidean distances and average linkage with the R function hclust. Based on the clustering, sample PUN7_LD1_S2_R2 was removed as an outlier (Fig. S1). To visualize major expression patterns and detect potential outliers among samples, principal component analysis (PCA) was performed on the expression matrix using the R function prcomp.

To estimate the contribution of experimental factors to gene expression variation (Oomen & Hutchings, 2022), we used the R packages variancePartition (version 1.38.1) and DESeq2 (Love *et al*., 2014). Variance components were estimated from the count matrix for each gene using linear mixed-effects models with sampling, room, and population treated as random effects, including all pairwise and three-way interactions. The proportion of variance attributable to each factor was calculated for each gene and averaged across genes to summarize the overall variance structure.

### Co-expression network analysis

To investigate the co-expression dynamics in the experiment, we performed a weighted co-expression network analysis using the WGCNA R package (Langfelder & Horvath, 2008). Co-expression network was calculated from the count matrix using the BlockwiseModules function with the options power = 8 (R^2^ > 0.9), mergeCutHeight = 0.25, minModuleSize = 30, reassignTreshold = 0, maxBlockSize = nGenes (24662), TOMType = “signed” and networkType = “signed”.

To evaluate associations between module eigengenes (MEs) and experimental factors, we fitted linear models in R with the function lm. Sampling time, treatment and population were included as fixed effects, along with all interaction terms. To account for multiple testing, the p-values for each model term were adjusted using Benjamini–Hochberg (BH) false discovery rate (FDR) correction for multiple testing.

### Gene Ontology enrichment

Gene ontology annotations were obtained from PANZZER2 (Ord *et al*., unpublished) and used to construct custom gene-term mappings. The enrichment analysis was conducted using the enricher function in the R package clusterProfiler (version 4.16.0) (Yu *et al*., 2012). The genes in each module were tested against a background consisting of all GO annotated genes (n = 21905). Enriched GO terms were identified using BH adjusted significance thresholds of p < 0.05 and q < 0.10.

### Differential gene expression analysis

All differential gene expression analyses were performed on the non-transformed count matrix in R (version 4.5.2) using the package DESeq2 (Love *et al*., 2014). We constructed a full interaction model including population, treatment, sampling time and all corresponding pairwise and three-way interaction terms to find differentially expressed genes between samples in different populations in different treatments and at different sampling time points. 951 genes were excluded due to non-convergence and the final number of genes included in the analysis was 23711.

Likelihood ratio tests (LRT) were used to evaluate the influence of individual factors and interaction complexity on model fit. In addition to the full model, we fitted a grouping model that combined all factors into a single composite variable to enable direct contrasts among specific combinations of population, treatment, and time combinations (e.g., differences between two populations under the LD1 treatment at sampling time 2).

### Circadian clock and flowering time genes

The *B. pendula* homologs for circadian clock and flowering time genes were identified from the reference proteome by DIAMOND (Buchfink *et al*., 2021) and BLASTP searches using BLAST+ v2.17.0 (Camacho *et al*., 2009). Protein sequences of circadian clock and flowering time regulators from *Arabidopsis thaliana* were retrieved from UniProt and used as queries. *B. pendula* homologs were identified by reciprocal best-hit searches using DIAMOND. To further identify FT-like genes, additional FT1 and FT2 protein sequences from *Populus tremula* were used as queries with BLASTP against the *B. pendula* reference proteome. Candidate homologs were assigned based on the highest-scoring matches and evaluated using PANZZER2 functional annotations (Table S2).

## Results

### Seedling growth patterns

We fitted a linear mixed model to analyse seedling absolute growth rate (mm/day) and separate the effects of population origin from the other factors (Fig. 2, Table 1, Fig. S2). Seedling growth rate was significantly influenced by sampling time, photoperiod treatment and population. Overall, the model effects were small (e.g., population effects ranging from −0.1 to 0.1 mm/day) as is expected with the short time scale of the experiment (Fig. S3).

**Fig. 2.**
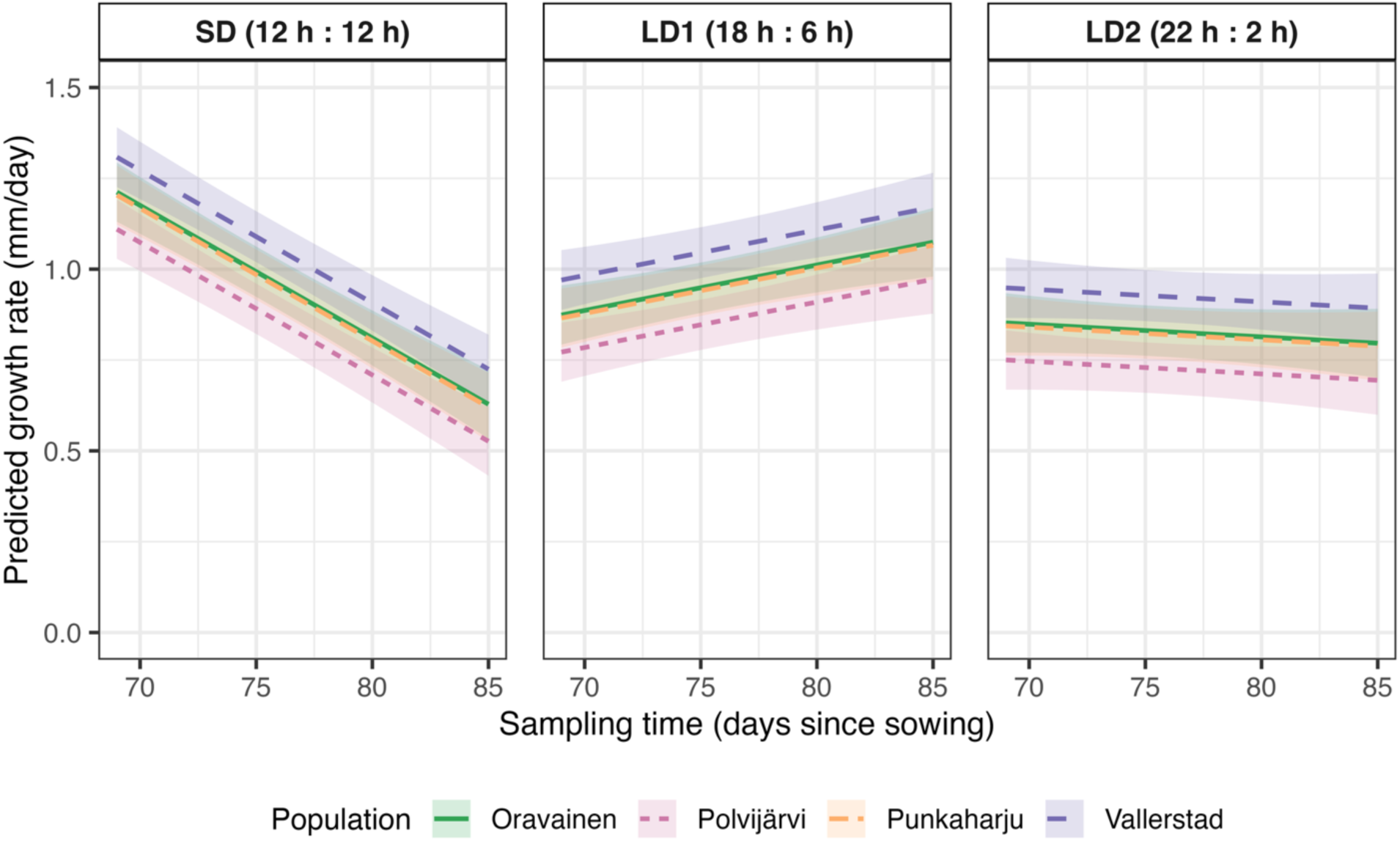
Predicted growth rates (mm/day) for *Betula pendula* seedlings as a function of sampling time for each population under different photoperiod treatments. Lines represent estimated marginal means derived from the fitted interaction model, and shaded ribbons indicate 95% confidence intervals. Panels are faceted by photoperiod treatment (SD 12 h : 12 h, LD1 18 h : 6 h, LD2 22 h : 2 h, hours light : dark), colours denote population, x-axis represents sampling time (days since sowing) and y-axis shows the predicted growth rates for each population.

**Table 1.** Fixed and random effects from the linear mixed-effect model of growth rate (mm/day) of *Betula pendula* seedlings under different photoperiods.

| Fixed effects: | Estimate | SE | df | t-value | p (>\|t\|) |
| --- | --- | --- | --- | --- | --- |
| *Intercept* | 0.977 | 0.034 | 46 | 28.392 | **< 2.00E-16** |
| *Sampling time* | -0.036 | 0.003 | 1853 | -10.476 | **< 2.00E-16** |
| *LD1* | -0.023 | 0.029 | 1853 | -0.800 | 0.4236 |
| *LD2* | -0.148 | 0.029 | 1853 | -5.110 | **3.55E-07** |
| *Population Polvijärvi* | -0.102 | 0.043 | 27 | -2.393 | **0.024** |
| *Population Punkaharju* | -0.008 | 0.043 | 27 | -0.191 | 0.850 |
| *Population Vallerstad* | 0.096 | 0.043 | 28 | 2.241 | **0.033** |
| *Sampling time × LD1* | 0.049 | 0.005 | 1853 | 9.963 | **< 2.00E-16** |
| *Sampling time × LD2* | 0.033 | 0.005 | 1854 | 6.682 | **3.11E-11** |
| **Random effects** | **Variance** | **SD** | **Number of groups** |  |  |
| *Mother tree (Intercept)* | 0.003 | 0.054 | 32 |  |  |
| *Residual* | 0.263 | 0.512 |  |  |  |

SE, standard error; df, degrees of freedom, SD; standard deviation. Statistically significant p-values in bold text.

In the short-day conditions, growth rate reduced with increasing duration of the photoperiod treatment (β = −0.036 ± 0.003, t = −10.48, p < 0.001). LD2 treatment reduced the growth rate further (LD2: β = −1.48 ± 0.029, t = −5.11, p < 0.001) while LD1 treatment did not have a significant effect (p < 0.42) compared to the SD treatment. However, in comparison to SD, the growth rate significantly increased over time in both long day conditions, (sampling time × LD1: β = 0.049 ± 0.005, t = 9.96, p < 0.001; sampling time × LD2: β = 0.033 ± 0.005, t = 6.68, p < 0.001). The increase in growth rate was significantly larger in LD1 compared to LD2 (post-hoc test; LD1 - LD2: β = 0.016 ± 0.005, t-ratio = 3.249, p = 0.0034).

Population effects were small (on average 0.07 mm/day), but mostly significant. Compared to the Oravainen population, growth rate was significantly reduced in the Polvijärvi population (β = −0.102 ± 0.043, t = −2.39, p = 0.024 and significantly increased in Vallerstad population (β = 0.096 ± 0.043, t = 2.24, p = 0.033). The growth rate in the Punkaharju population did not differ from Oravainen (p = 0.85), and these effects were not significantly influenced by sampling time or treatment effects as indicated by the worsened model fit when including interaction terms for population effects (Table S3).

Variance attributable to the random effect of the mother tree was low (σ^2^ = 0.003, SD = 0.054), whereas residual variance was greater (σ^2^ = 0.263, SD = 0.512), indicating that individual variation in growth rates was substantial even in controlled growth chamber conditions.

### Read quality, mapping, gene counts and exploratory analyses

In total, the dataset comprised 3.73 Tbp of raw 150 bp paired end (PE) sequencing data, corresponding to an average of 24.93 million PE reads per sample (Table S1). After quality filtering and trimming of adapters, on average 23.45 million PE reads per sample remained. The average mapping rate to the reference genome was 96.29 %, with on average 93.65 % of reads mapping uniquely. On average only 56.24 % of reads mapped uniquely to the tetraploid *B. pubescens* genome and no outliers were observed, indicating that the samples are *B. pendula* (Fig. S4).

We generated a combined gene-level count matrix across all samples comprising 34319 genes and 214 samples using featureCounts. Across samples, the majority of mapped fragments were assigned to genes, with assignment rates ranging from 86.33 – 94.09 % while most of the remaining fragments (6.31 %) were predominantly unassigned due to lack of overlap with annotated exons. A small fraction (1.34 %) was ambiguous corresponding to reads overlapping multiple genes or exons. After low-count filtering, the final expression matrix comprised 24662 genes.

The PCA did not show clear clustering of samples to treatments, sampling times or populations (Fig. 3), indicating that the major axes of variation in gene expression were dominated by individual variation.

**Fig. 3.**
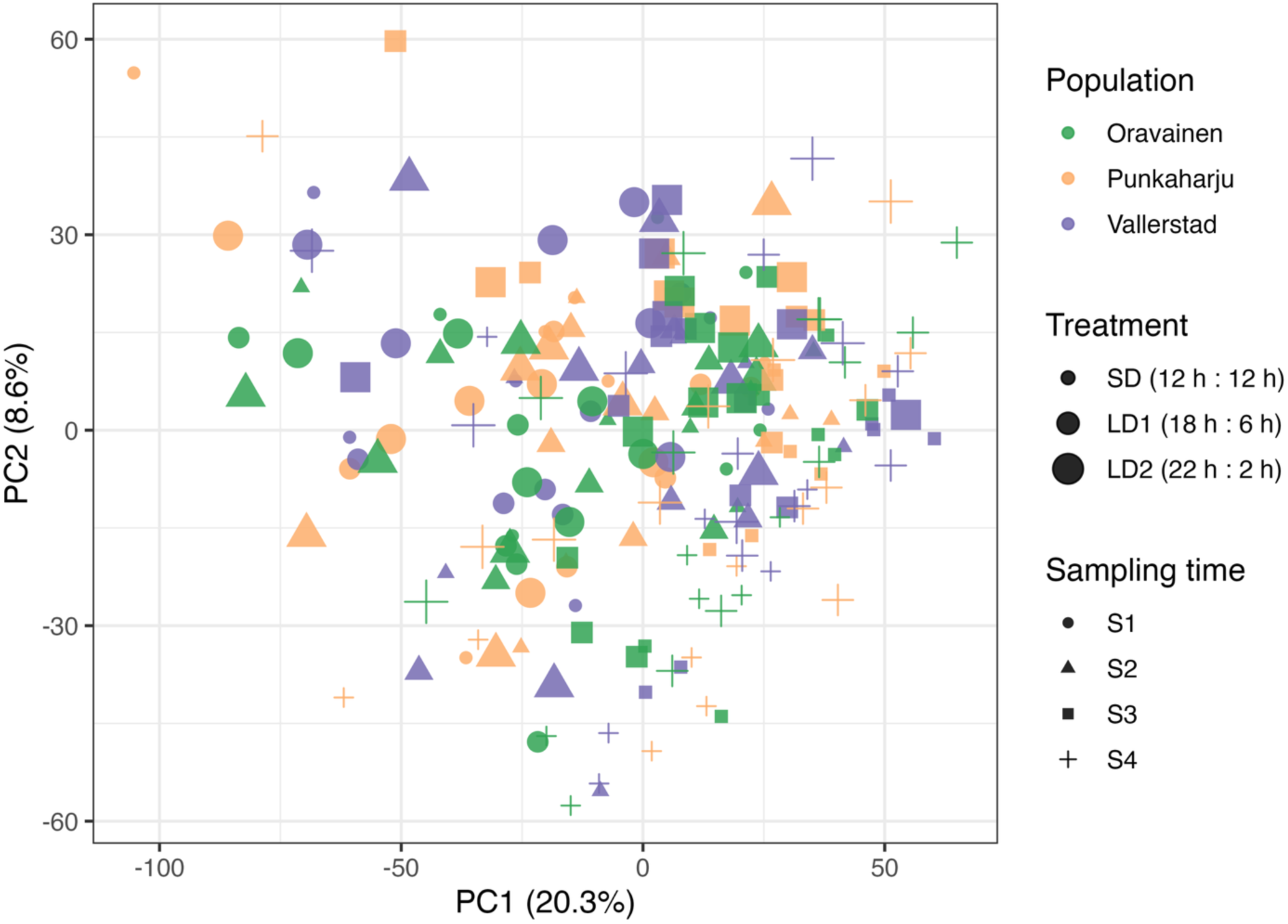
Principal component analysis (PCA) of variance-stabilized gene expression data from *Betula pendula* seedlings. Each point represents a sample, positioned according to the first two principal components (PC1, PC2).

### Variance partitioning

To identify population and whole transcriptome level differences in gene expression, and interactions with time and treatment, we employed variance partitioning, weighted co-expression network analysis and GO term analysis. Consistent with the PCA, variance partitioning indicated that residual variance accounted for 82.26 % of total gene expression variation (mean across genes), reflecting large individual variation across samples (Table 2). Of the modelled terms, sampling time explained the largest proportion of variance (6.89 %), followed by the sampling time × treatment interaction (5.64 %) and treatment (2.38 %). Population explained 1.52 %, treatment × population interactions 0.46 % and sampling time × population interactions less than 0.01 % of the variance. The three-way interactions explained 0.84 % of the variance which is a larger fraction than the 2-way population-related interactions indicating that population-specific responses often depend on both sampling time and treatment.

**Table 2.** Variance partition for variance stabilized gene expression data from *Betula pendula* seedlings with mean variance explained by the modelled factors across all genes.

| Variance explained | % |
| --- | --- |
| Residuals | 82.26 |
| Sampling time | 6.89 |
| Sampling time × treatment | 5.64 |
| Treatment | 2.38 |
| Population | 1.52 |
| Sampling time × treatment × population | 0.84 |
| Treatment × population | 0.46 |
| Sampling time × population | < 0.01 |

Sampling time was the factor explaining most of the expression variance for 142 genes, treatment for 38 genes, sampling time × treatment for 20 genes, and population for one gene (Table S4, Fig. 4).

**Fig. 4.**
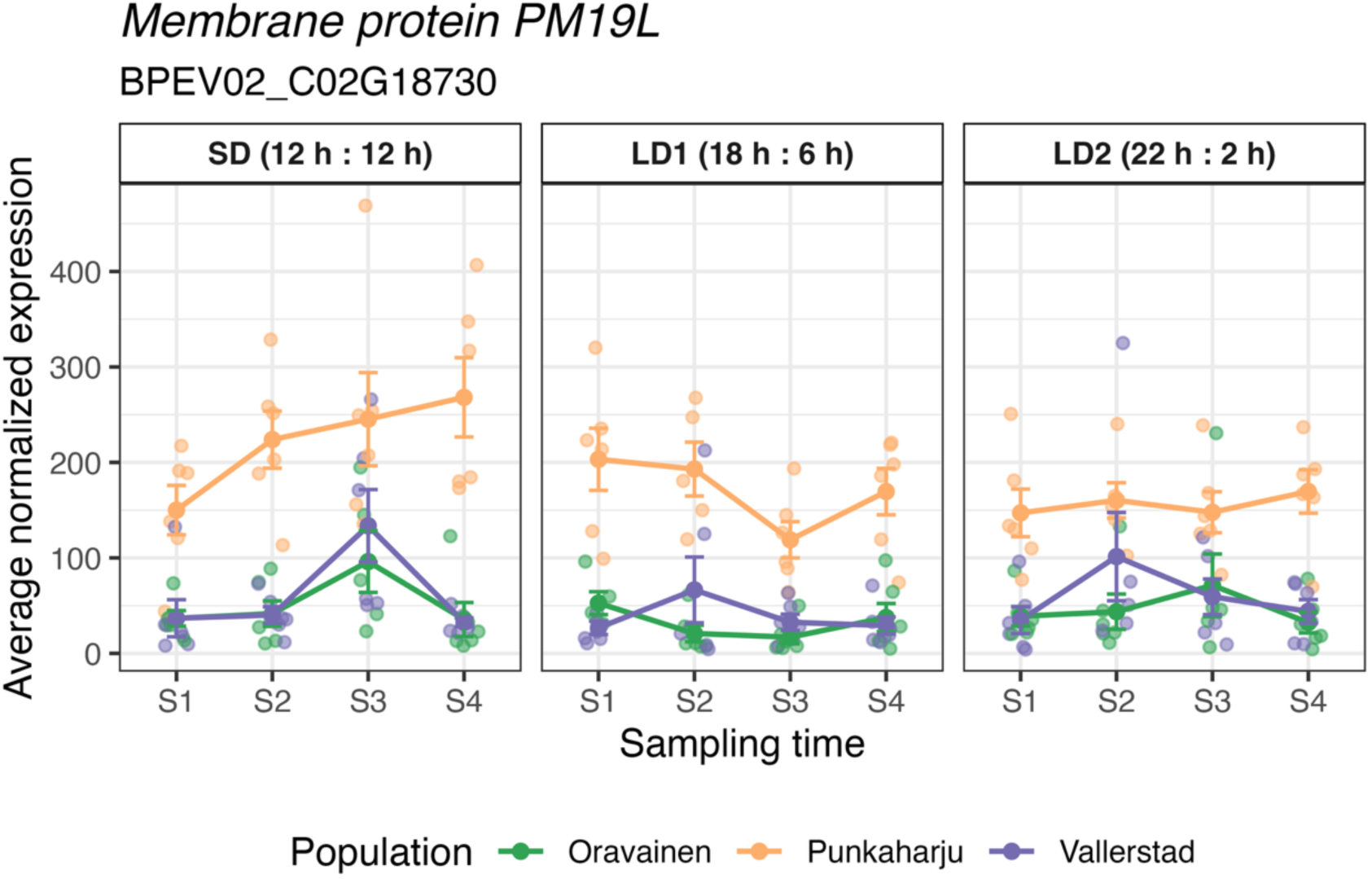
Expression of *PM19L* for which population explains most variance in expression in *Betula pendula* seedlings in different photoperiod treatments. Lines indicate population averages in expression with vertical bars indicating standard error in expression within populations.

### Weighted co-expression network analysis

To study the level of functional pleiotropy across sampling time points, treatments, and populations, we performed weighted gene co-expression network analysis (WGCNA). The analysis produced 20 co-expression modules with a total of 17408 (70.59 %) of genes assigned to modules (Table S5, module genes listed). The remaining 7254 (29.41 %) genes remained unassigned (module grey). Given that 66.78 % of the total 34319 genes in the *B. pendula* genome are annotated, genes in co-expression modules were enriched for annotated genes (Table 3). Based on the annotation and GO enrichment, most modules form logical functional units, e.g., cell cycle (blue), translation (brown), photosynthesis (magenta), circadian rhythm (cyan), and defence and immune response (turquoise). Module grey, that represents genes not assigned to any co-expression module, had the lowest number of annotated genes (63.5 %). In contrast, module grey60 (vacuolar targeting and transcriptional regulation) had the most annotated genes (95.95 %) followed by modules red (chloroplast organization, 95.35 %) and cyan (circadian rhythm and photoperiodic regulation, 95.33 %). Due to specificity of the Biological Process (BP) GO terms per each module (Fig. 5, Table S6), we named the modules based on their functions.

**Fig. 5.**
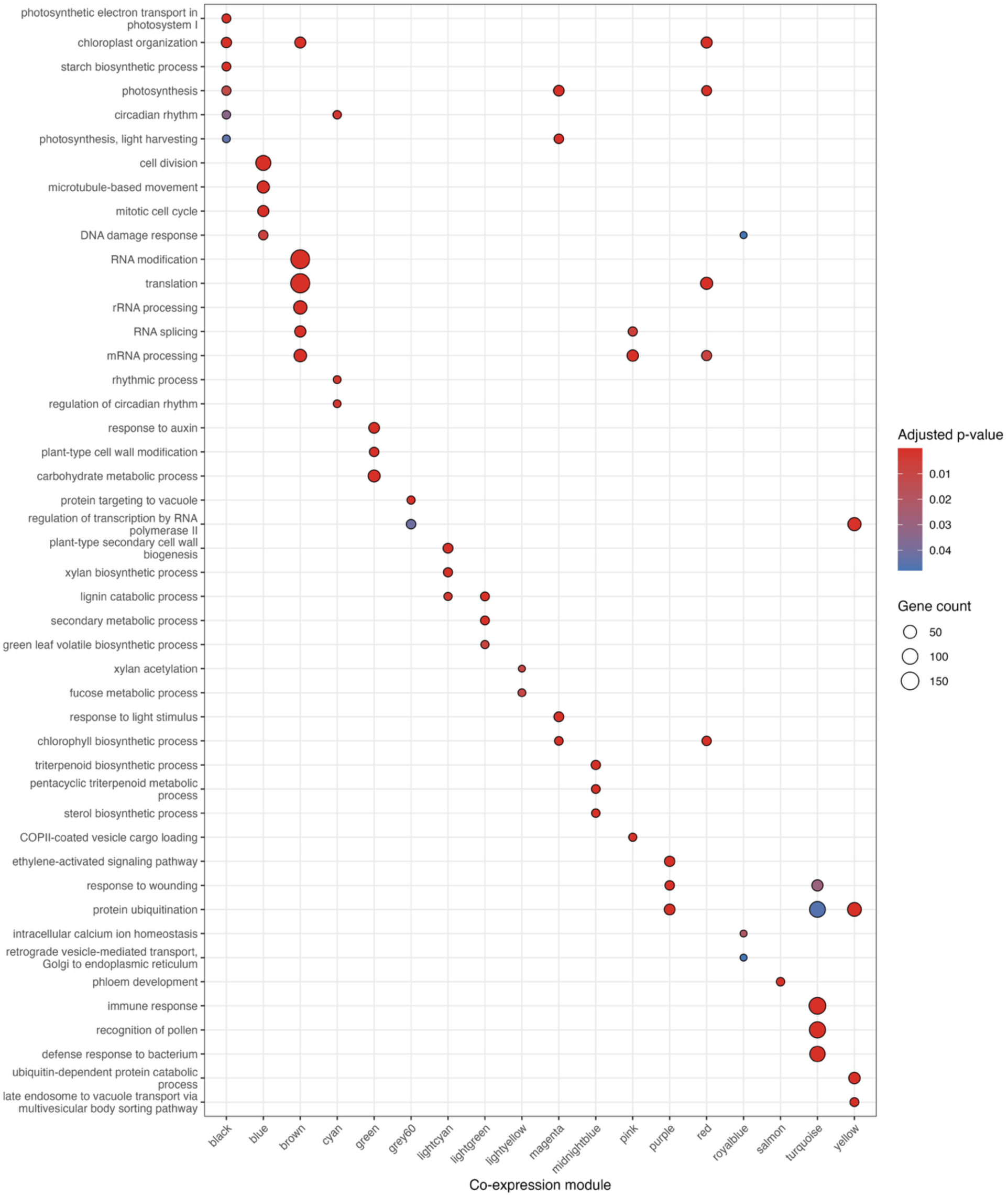
Gene Ontology (GO) enrichment of the weighted gene co-expression network analysis (WGCNA) modules for biological processes (BP) in silver birch (*Betula pendula*) seedlings. Enrichment was tested separately for each module using compareCluster with the enricher function in clusterProfiler (Yu *et al*., 2012). P-values were adjusted with Benjamini-Hochberg false discovery rate correction. Tan and greenyellow modules did not have significant BP enrichment. The unassigned genes (grey module) were excluded from the enrichment analysis.

**Table 3.**
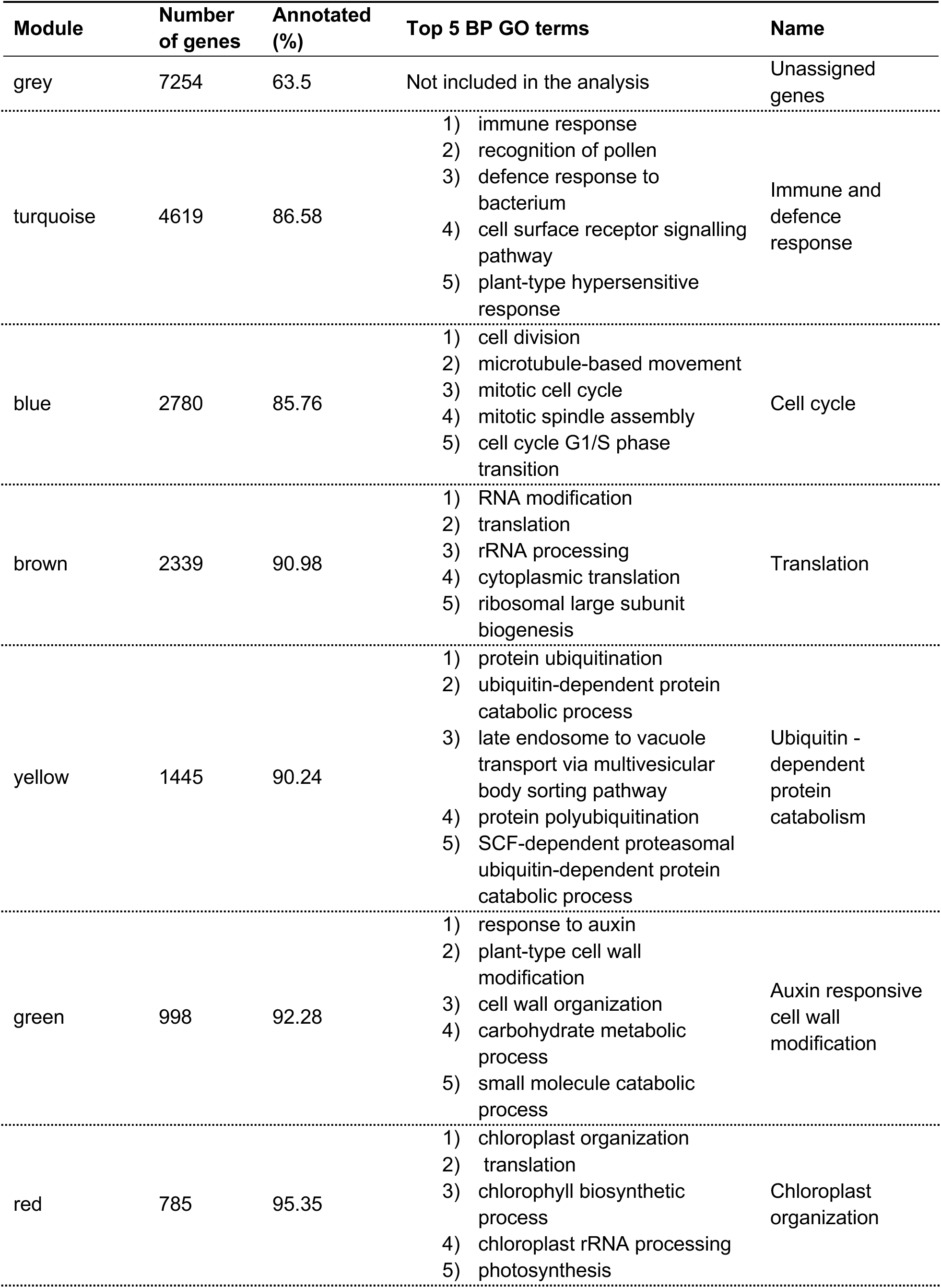
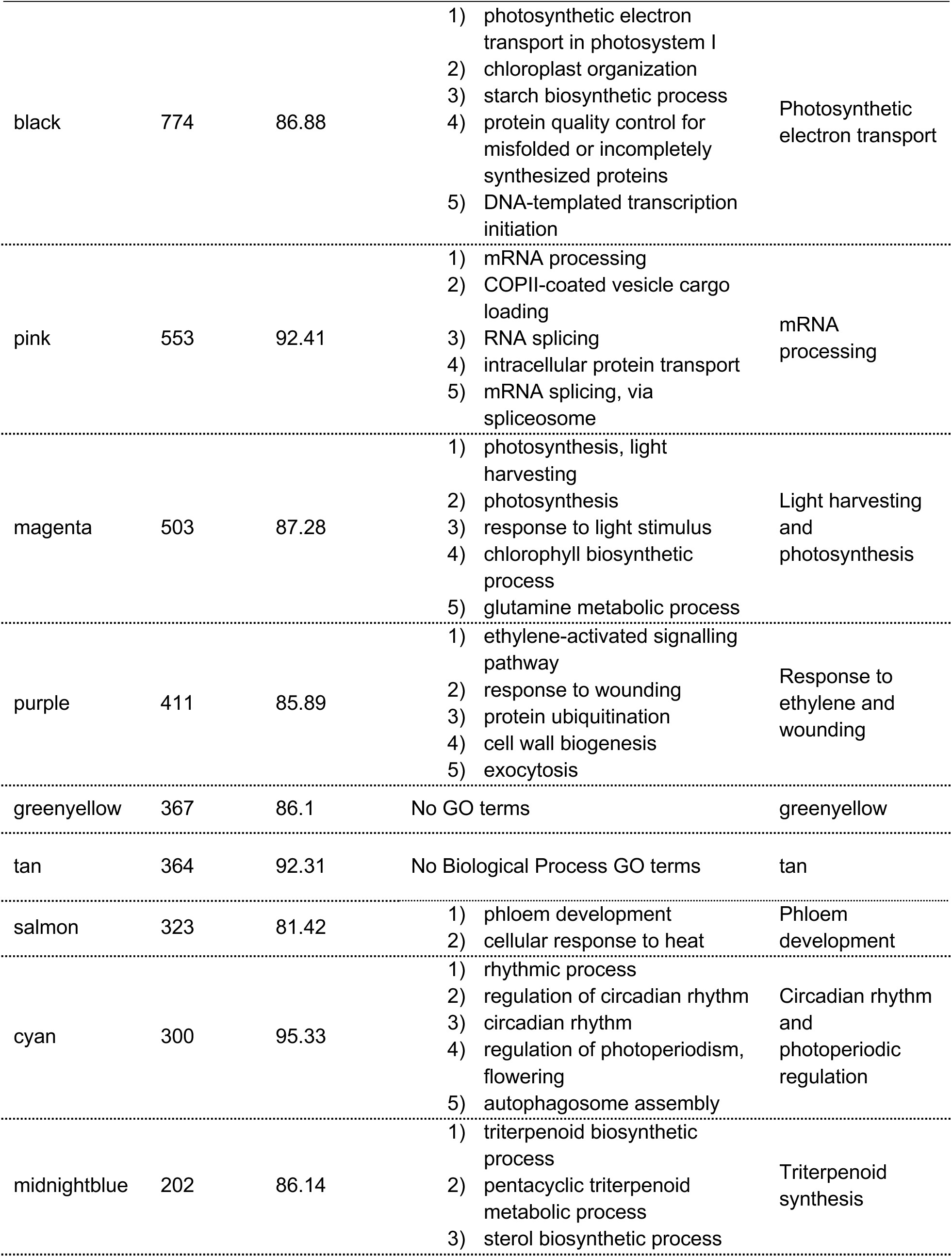
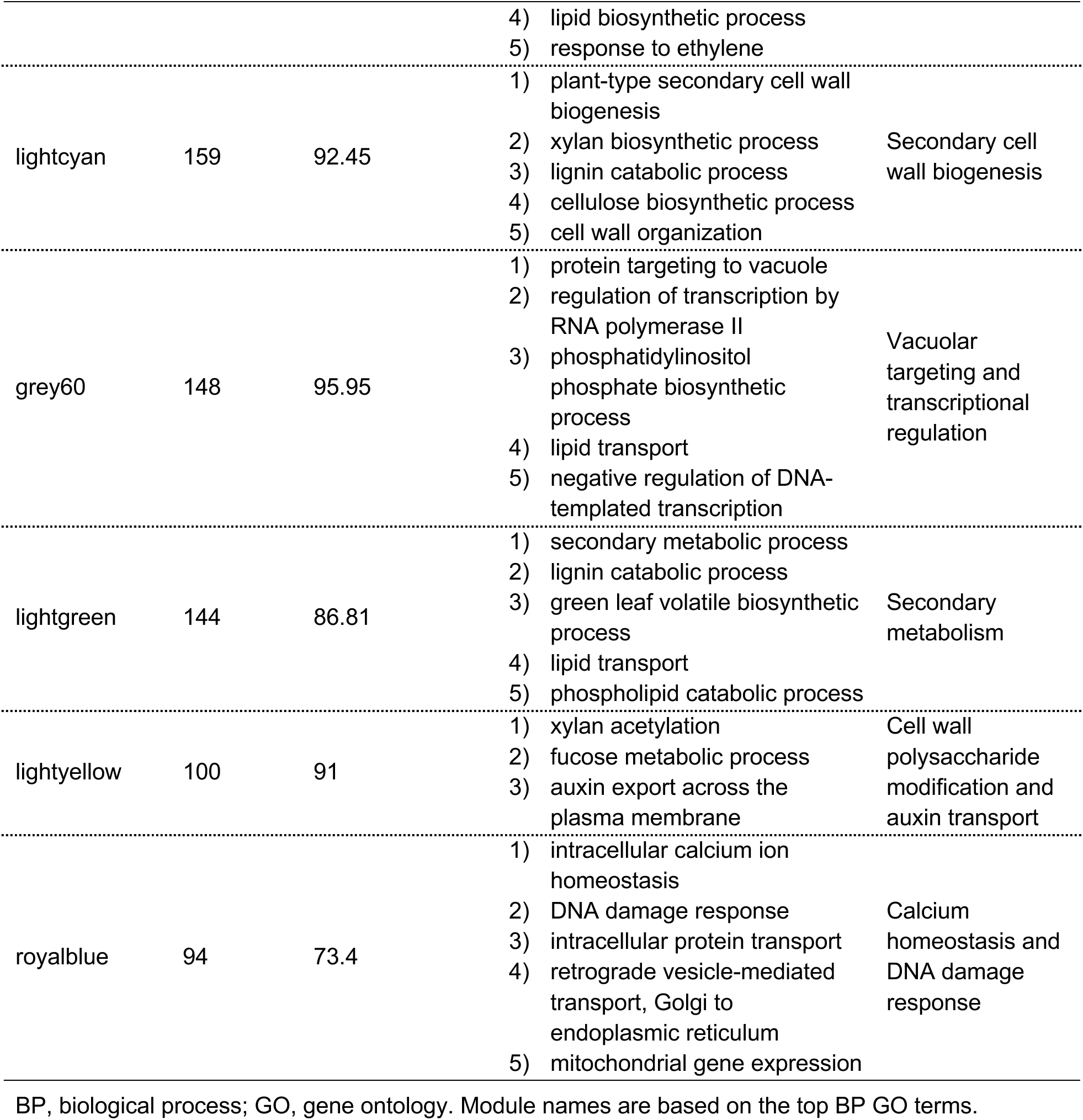
Summary table of co-expression modules produced by weighted gene co-expression network analysis (WGCNA) on *Betula pendula* seedlings annotated with Biological Process Gene Ontology terms.

When associating module eigengene (ME) expression with the treatments using linear models, sampling time and its interactions with treatments had the most associations (13 modules) (Fig. 6a). Notably, in the Oravainen baseline, sampling time in the very beginning of the photoperiod treatment (S2) induced significant expression changes only in two modules: circadian rhythm and photoperiod regulation (cyan) and calcium homeostasis and DNA damage response (royalblue) (Fig. 6a). Functionally wider impacts were evident at later sampling times, 3 and 9 days after photoperiod onset (S3 and S4, Fig. 6a). Similar trends were evident also in other population baselines (Fig. S5).

**Fig. 6.**
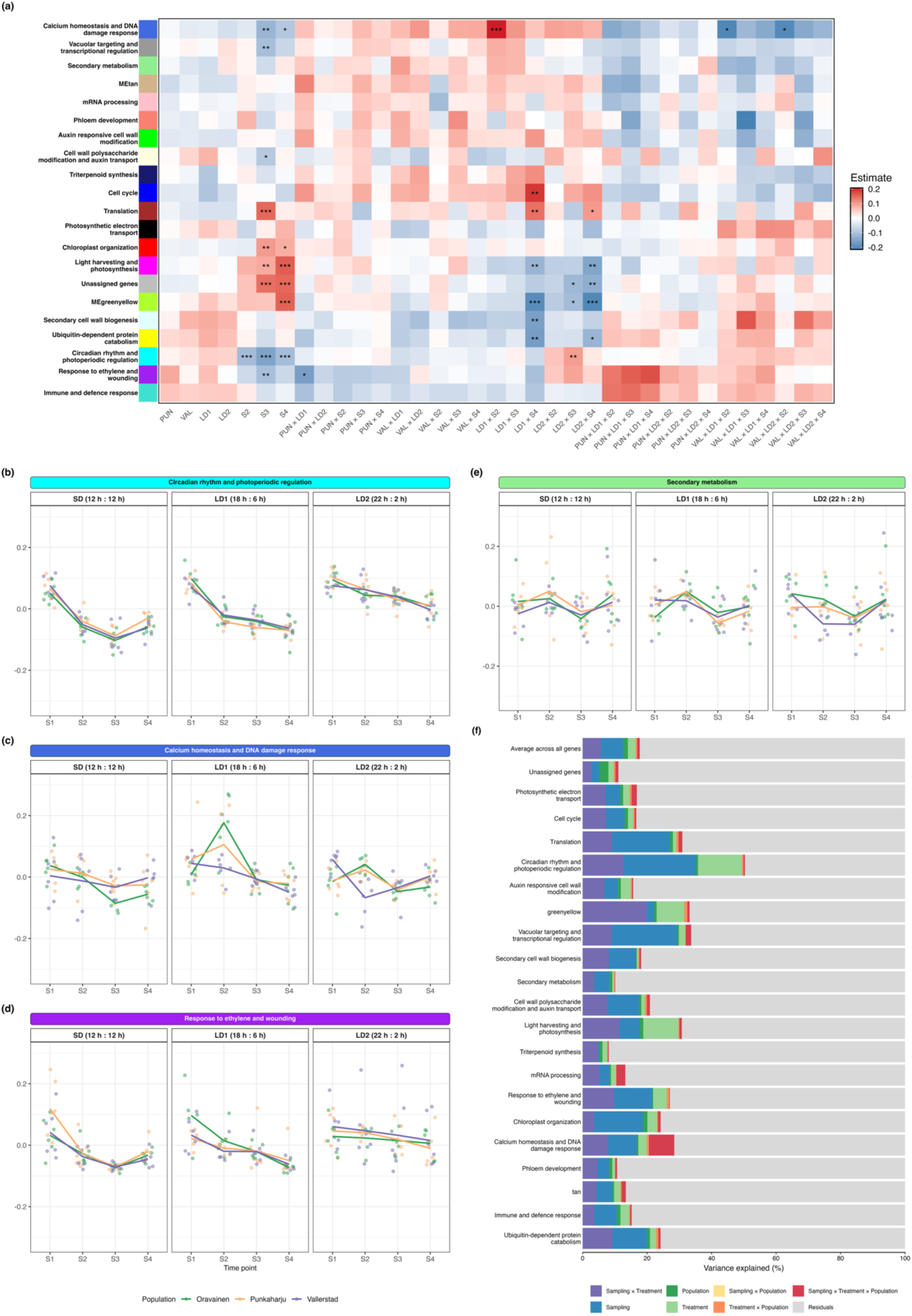
(a) Relationship of the co-expression module eigengene expression to the treatments in *Betula pendula* seedlings. Heatmap showing regression coefficients from linear models fitted to module eigengenes (MEs) of weighted gene co-expression network analysis (WGCNA) modules. Oravainen population at sampling time 1 in constant light (SD room) was used as the baseline. Each cell of the heatmap represents the estimated effect of a model term (x-axis) on a given module eigengene (y-axis). Colour indicates the direction and magnitude of the effect (blue, negative; white, no effect; red, positive). Asterisks denote false discovery rate adjusted significance levels (**\*\*\***: *p* ≤ 0.001, **\*\***: *p* ≤ 0.01, **\***: *p* ≤ 0.05). (b-e) Selected module eigengene expression across populations and treatments. Lines indicate the mean eigengene value for each population across sampling time points. Panels are faceted by treatment. Each subplot corresponds to a single co-expression module from WGCNA. All modules in Fig. S6. (f) Variance components for WGCNA modules. Variance components averaged over all genes in each module. The topmost bar indicates the average over all genes. Colours indicate the different variance components related to the experimental design.

The impacts of photoperiod treatments were primarily time dependent. For instance, the circadian rhythm and photoperiod regulation (cyan) ME expression generally decreased with time, but the rate of decrease depended on the treatment (Fig. 6b). The ME expression decreases slower in the longest day (LD2) treatment in comparison to SD and LD1 treatment. The ME expression was consistent among populations and variation was low between samples, indicating a coordinated response to photoperiod change from S1.

Population effects in ME expression were less prevalent than sampling time and photoperiod treatment effects. Two modules showed significant differences in ME expression between populations: calcium homeostasis and DNA damage response (royalblue, Fig. 6c) and response to ethylene and wounding (purple, Fig. 6d). Response to ethylene and wounding ME expression was significantly lower in the LD1 treatment in the Punkaharju population compared to the Oravainen population. Calcium homeostasis and DNA damage response ME expression was significantly lower in the Vallerstad population compared to the Oravainen population at sampling time 2 in both long day treatments. In LD1, the ME expression was significantly higher in Oravainen at sampling point 2 compared to the SD treatment.

In total, five modules (secondary metabolism (lightgreen), tan, mRNA processing (pink), auxin responsive cell wall modification (green), and triterpenoid synthesis (midnightblue) were not associated with any experimental factor in any population baseline, indicating a large group of functions whose co-expression is controlled by other factors than sampling time, photoperiod and population (Fig. S5, Fig. S6, example module in Fig 6e).

Variance partition for module genes (Fig. 6f) indicated that expression variation in the circadian rhythm and photoperiod regulation module was to a large extent explained by the photoperiod, sampling time and their interactions. Notably, the calcium homeostasis and DNA damage response module’s expression stood out as explained by the population × sampling time × treatment interactions clearly more than any other module (7.9%, Fig. 6f, Table S7).

### Differential gene expression

For individual gene level analysis, we employed a full interaction model and tested the significance of all three factors and their interactions with LRTs. Sampling time, photoperiod treatment, and their two-way interactions had the largest influence on gene expression, affecting 15199, 10459, and 6125 genes respectively (Table 4, genes listed in Table S8). 1589 genes (6.7 %) had population-level differences in gene expression. Far fewer genes (172 genes, 0.73 %) were influenced by population × treatment interactions and population × sampling time interactions (161 genes, 0.68 %). 28 genes (0.12 %) showed a three-way interaction, indicating that a minor fraction of the genes exhibited differences between populations in a certain treatment at a specific time point.

**Table 4.** Likelihood ratio test (LRT) for different factors included in the differential gene expression model with the number of genes that were significantly associated with each model term.

| Factor tested | Number of significant genes | % of significant genes | % of unassigned genes |
| --- | --- | --- | --- |
| Sampling time | 15199 | 64.11 | 15.7 |
| Treatment | 10459 | 44.12 | 18.0 |
| Population | 1589 | 6.7 | 48.1 |
| Any interaction | 6012 | 25.36 | 17.1 |
| Any population interaction | 376 | 1.59 | 35.6 |
| Population × sampling time | 161 | 0.68 | 46.0 |
| Population × treatment | 172 | 0.73 | 36.0 |
| Sampling time × treatment | 6125 | 25.84 | 15.6 |
| Population × sampling time × treatment | 28 | 0.12 | 64.2 |

The majority (64.2%) of the genes for which the 3-way interactions improved the model fit were not co-expressed in any module (Table 4). Further, 48.1 % of the genes with significant population effects and 35 % of the genes involved in population interactions were not co-expressed. In contrast, only 15.7 % and 18 % of sampling time and treatment genes, respectively, were not co-expressed. This indicates that genes responding to sampling time and treatment and their interactions are more likely to be co-expressed than genes whose expression largely depends on population or population interactions with sampling time and treatment. Overall, genes whose expression depended on population effects were 2.23 times more likely to be unassigned to co-expression modules than expected (Table S9).

Based on the likelihood ratio test results (Table 4) and our *a priori* specific objective to identify population-specific time-sensitive photoperiod responses, we specified a full interaction model for DGE analysis (Fig. 7). For each DEG with module membership, we also obtained its connectivity metric (kME; correlation to the module eigengene expression, Table S10). In pairwise contrasts, largest differences were found between sampling times (S3 vs S1; 3318 DEGs, S4 vs S1; 3069 DEGs) and sampling time-treatment interactions, especially at sampling time 4 in LD1 treatment (4193 DEGs) and at sampling time 4 in LD2 treatment (1637 DEGs). Notably, when averaged over populations and sampling time points, LD1 treatment induced more differences (573 DEGs) than longer day LD2 treatment (36 DEGs) when compared to the baseline (Oravainen in constant light (SD treatment room) at sampling time 1).

**Fig. 7.**
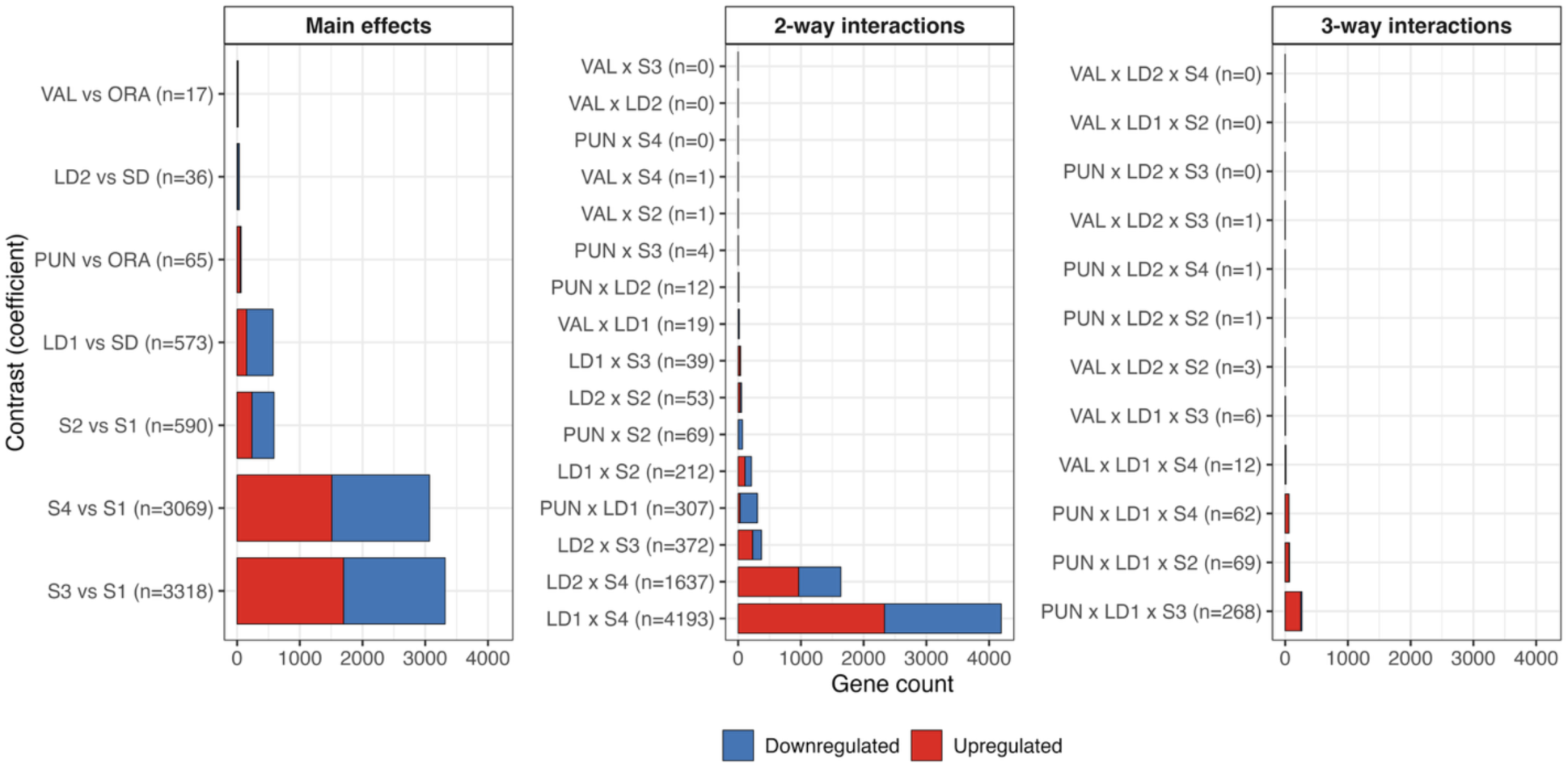
Summary of differentially expressed genes (DEGs). Bar plots summarizing the number of DEGs detected for each coefficient of the differential expression model. For each contrast, red bars represent the number of upregulated genes and blue bars downregulated genes relative to the reference condition (population Oravainen in constant light (SD treatment room) at sampling time 1). Significance is based on an adjusted p-value threshold (padj < 0.05) and an absolute log₂ fold-change > 0.

Averaged over time points and treatments, there were 65 DEGs between Punkaharju and Oravainen and 17 DEGs between Vallerstad and Oravainen that did not depend on sampling time or treatment. Gene expression between Vallerstad and Oravainen was consistently very similar throughout the experiment with at most 19 DEGs between the populations (VAL × LD1).

The overall effect of sampling time to population differences averaged over treatments was not prominent - the largest difference was between Punkaharju and Oravainen at sampling time 2 with 69 DEGs (PUN × S2), most of them downregulated in Oravainen. The largest differences between populations were between Punkaharju and Oravainen in the LD1 treatment (307 DEGs). When looking at specific time points, sampling time 2 was associated with 69 DEGs, sampling time 3 with 268 DEGs and sampling time 4 with 62 DEGs between Punkaharju and Oravainen. LD2 treatment was associated with at most 12 DEGs between the populations when averaged over time points.

Number of DEGs, module membership and average connectivity of the DEGs (Fig. 7, Table S10) indicated unique and coordinated functional response (Immune and defence response, Response to ethylene and wounding) in the Punkaharju population in the LD1 treatment consistently throughout all sampling points. In other contrasts, most between-population DEGs were unassigned (50-100 %).

### Circadian clock and flowering time gene expression patterns

We studied the expression patterns of circadian clock and flowering time genes in more detail due to their central role in photoperiod response and adaptive variation. The mean variance partitioning across homologs the circadian clock and flowering time genes indicated stronger than average sampling time, treatment and sampling time × treatment interaction effects than on average across all genes (Fig. 8a), but no outstanding population level effects. However, some individual genes’ variance partitioning exhibited also population and population interaction effects (e.g., *APRR1/TOC1* (BPEV02_C12G09180), *CONSTANS-like* genes (BPEV02_C02G18100, BPEV02_C10G01710), *CDF1* (BPEV02_C07G18430).

**Fig. 8.**
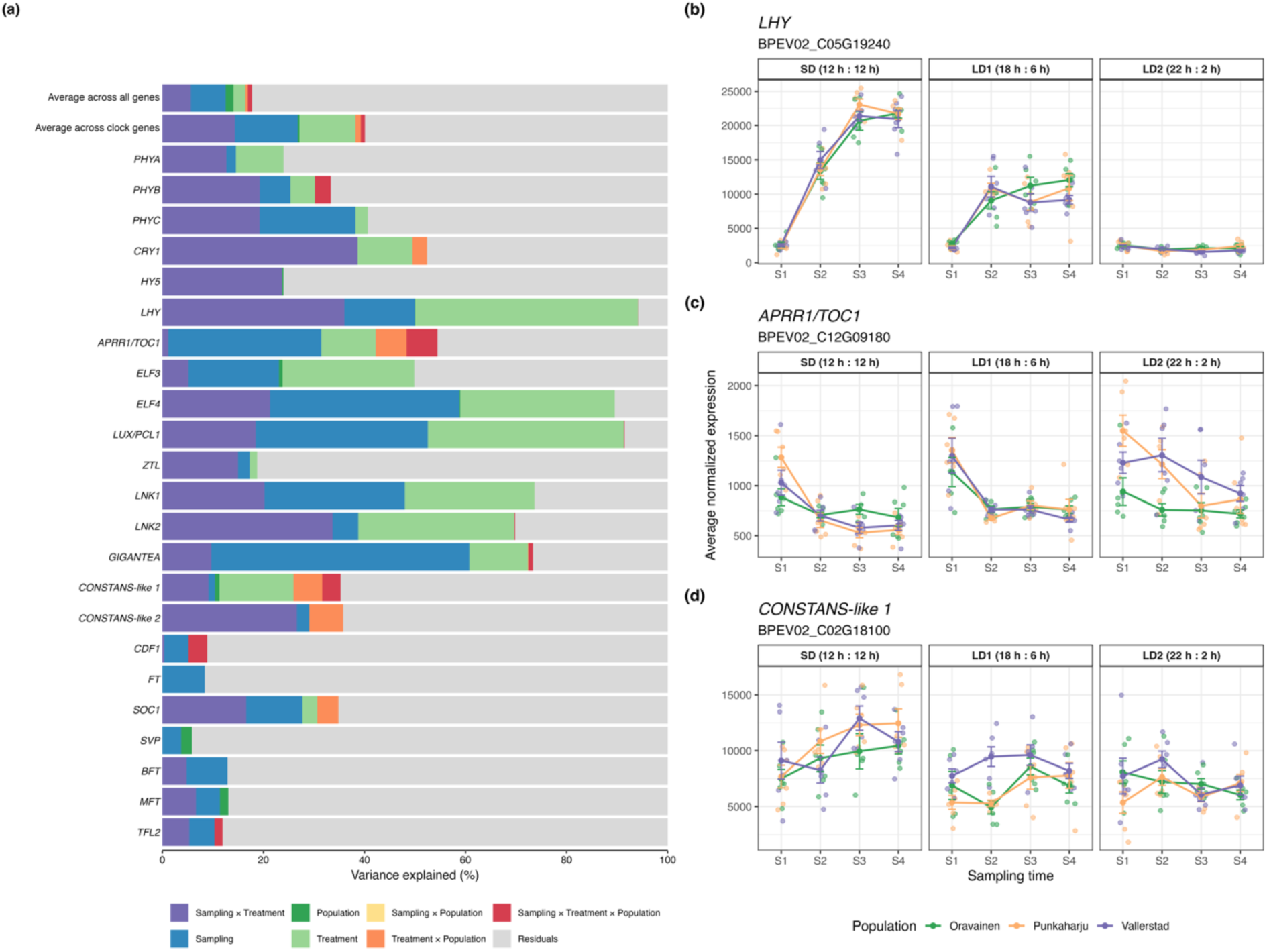
(a) Variance partition for selected circadian clock and flowering time genes in *Betula pendula* seedlings. Includes average over all genes used in the analysis (n = 24662) and average of the shown genes. Expression plots for all the circadian clock and flowering time genes included in this study in Fig. S7. (b-d) Expression of selected circadian clock genes. Lines indicate population averages in expression with vertical bars indicating one standard error in expression within populations.

*LHY/CCA1* (BPEV02_C05G19240) showed one of the most prominent effects of photoperiod treatment with a drastic expression increase in SD and LD1 across populations (padj = 2.22E-130, sampling x treatment LRT test, Table S8, Fig. 8b), whereas population impacted the expression of e.g., *APRR1/TOC1* (Fig. 8c) and *CONSTANS-like 1* (BPEV02_C02G18100) (Fig. 8d)*. APRR1/TOC1* expression was higher in Punkaharju and Vallerstad populations compared to Oravainen in the LD2 treatment. As indicated by the variance partition, the expression of *CONSTANS-like 1* depended on the population × treatment effect as the expression was higher in the Vallerstad population compared to the others in LD1. This effect was significant when analysing the time points separately but not in the LRT (Table S8).

From the 24 circadian clock and flowering time genes (Fig. S7), three (*ELF4* (BPEV02_C05G01780), *GIGANTEA* (BPEV02_C06G07910), *LUX/PCL1*(BPEV02_C09G05890) were found in the circadian rhythm and photoperiodic regulation module where their expression correlated strongly with the ME expression (kME) and they were highly connected (kWithin) with other genes in the module (Table S11, kME: 0.75-0.86, kWithin: 47.9-67.8, top 75 connectivity percentile). This modules’ expression decreased with time during the experiment. *LHY/CCA1* and *CRY1* (BPEV02_C10G13050), whose expression increased with time especially in SD and LD1 treatments, were placed in the greenyellow module (kME = 0.66, kWithin = 41.7 and kME = 0.84, kWithin = 52.8 respectively, top 90 connectivity percentile).

## Discussion

Changes in the regulation of gene expression have been shown to mediate adaptive phenotypic differences across complex traits (Ayroles *et al*., 2009), leading us to hypothesize that locally adapted *B. pendula* populations would show differences also in transcriptomic phenotypes. We used gene expression patterns to study transcriptome wide population level differences in *B. pendula* first year seedlings. Seedlings were subjected to photoperiod treatments to trigger differences in photoperiod response quantity and quality among populations (Taulavuori *et al*., 2017). The sampled populations show low genetic differentiation (FST < 0.01, Milesi *et al*., 2024; Herrera-Egoavil *et al*., 2026) and represent a fraction of complete latitudinal and environmental clines that likely extend across the species range. The benefit of studying tree populations in fairly close geographic proximity is that results are unlikely to be affected extensively by population structure and population-specific drift; rather, phenotypic differences are to a large extent expected to be maintained by natural selection.

### Growth differences among populations are subtle but indicate local adaptation and expected responses to photoperiod changes

Focusing on geographically relatively proximate populations and early transcriptomic photoperiod responses allowed us to pinpoint previously unknown patterns of local adaptation in *B. pendula*. Differences in growth among populations, despite being small, were in line with earlier studies from common gardens showing clinal patterns in phenological and growth traits (Li *et al*., 2003; Viherä-Aarnio *et al*., 2005; Tenkanen *et al*., 2023) indicative of local adaptation. The most southern Vallerstad population grew fastest, but the three other populations did not follow a latitudinal pattern in their growth. The difference in latitude among Oravainen, Punkaharju and Polvijärvi were very small (1.55 degrees) and may be overridden by continental-coastal patterns of climate. The observed reduction in growth rate as a response to SD treatment was expected as the short-day length is a known trigger for growth cessation in *B. pendula* (Li et al. 2003) and boreal forest trees in general (Singh *et al*., 2017). Unexpectedly, growth was slower under longer days (22 h, LD2) than under 18 h LD1 treatment. In natural conditions, assuming the initiation of seedling growth in the beginning of June (day length in Oravainen 19.5 h, Punkaharju 19 h, Vallerstad 17.8 h), the sampling time would correspond to the beginning of August (day length in Oravainen 17.5 h, Punkaharju 17.3 h, Vallerstad 16.5 h). Thus, the LD1 is closer to natural conditions of the populations which could explain the optimum growth under these conditions. Li *et al*., 2003 determined the critical daylength for growth cessation for populations with roughly the same latitude origins as Vallerstad and Oravainen populations to be 14 h and 18 h, respectively. This is congruent with our observation of significant reduction in growth at 12 h SD daylength in comparison to 18 h and 22 h in all studied populations.

### Gene expression changes are driven by photoperiod and time, with small but significant population impacts

All the analytical approaches (variance components, co-expression modules and DGE) indicate that sampling time has the largest impact on gene expression pattern, followed by photoperiod treatment and sampling time × treatment interactions. Differences in gene expression increased and interactions became more significant by time. For example, in comparison to baseline, LD1 × S4 had the most DEGs in the whole experiment. The pattern was expected as both developmental stage and response to treatment are time-dependent and photoperiod is expected to have a significant impact on gene expression. Despite this, significant gene expression differences among populations were also detected at whole-genome, module- and single gene level.

At the whole transcriptome level, population and its interaction effects explained on average less than 3 % of total gene expression variance. However, population played a significantly larger role in e.g., calcium homeostasis and DNA damage response (8.94 %) and mRNA processing (3.43 %) modules compared to the average of all genes (Fig. 6). These results can potentially be explained by calcium ions having a role in plant stress signalling in response to different biotic and abiotic stresses (Estravis-Barcala *et al*., 2020; Ren *et al*., 2023), and calcium-signalling related genes have been identified in local adaptation scan of *Abies religiosa* (Arenas *et al*., 2025) and responding to temperature stress in Sitka spruce (Holliday *et al*., 2008).

Single genes exhibiting population-dependent expression differences and assigned to co-expression modules were most frequently associated with immune response and defence, ethylene and wounding, as well as calcium homeostasis and DNA damage response modules (Table S10). A homolog of *PM19L* was identified as the singular gene for which population explains most of its expression variance (Fig. 4, Table S4). *PM19L* is a member of the AWPM-19 gene family which is characterized by a conserved function in drought tolerance both in *Arabidopsis* and *Populus* (Wang *et al*., 2022). *PM19L* homologs have been associated with tolerance to a variety of abiotic stresses in different species, such as drought and salinity resistance in peony (Meng *et al*., 2022) and osmotic stress in rice (Chen *et al*., 2015). In *Arabidopsis* and pearl millet, *PM19L* has been linked to seed germination under osmotic and salinity stress respectively (Yu *et al*., 2024; Alexander *et al*., 2025). Given the role of PM19L proteins in drought and osmotic stress tolerance the population specific expression pattern could contribute to local adaptation by influencing the capacity to maintain cellular water balance under different growing conditions.

Even though the exact functions and underlying molecular mechanisms of these genes in *B. pendula* remain uncharacterized, the genes with population specific responses are involved in various stress responses. This suggests that birch populations are adapted to not only photoperiod, but also other local environmental factors such as soil composition, moisture, inter-specific competition, herbivores and pathogens.

### Gene expression patterns in known circadian clock genes

Most evidence for population differentiation in photoperiodic responses in forest trees derives from candidate gene and QTL studies focusing on components of the photoperiodic and circadian clock pathways, including *FT*, *CONSTANS*, phytochromes and associated regulators (Ingvarsson *et al*., 2006; Böhlenius *et al*., 2006; Fabbrini *et al*., 2012; Avia *et al*., 2014; Ranade & García-Gil, 2023). By contrast, transcriptome-wide comparisons of photoperiodic responses among populations remain understudied, especially through time (but see Cañas *et al*., 2015; Hess *et al*., 2016).

Given the central role of circadian clock regulation in adaptation to seasonal variation in day length (Böhlenius *et al*., 2006), we expected to detect population-level differences in the expression of core circadian clock and flowering time genes. Instead, expression variation in most of these genes both on the individual and co-expression module level was driven predominantly by sampling time, photoperiod treatment, and their interaction (Fig. 6f, 8a). Indeed, some of the strongest transcriptional responses to the photoperiod treatment were observed among homologs of the circadian clock components, including *LHY/CCA1* (BPEV02_C05G19240), *ELF4* (BPEV02_C05G01780), *LNK1* (BPEV02_C03G11350), and *GIGANTEA* (BPEV02_C06G07910). For these genes population effects explained a negligible fraction of the variance.

Despite the small population effects in most of the circadian clock and flowering time genes, the homologs of a core circadian clock component *APRR1*/*TOC1* (BPEV02_C12G09180) and a flowering time regulator *CONSTANS-like 1* (BPEV02_C02G18100) were differently expressed between populations (Fig. 8b-c). The expression of *CONSTANS-like 1* in the Vallerstad population in LD1 at sampling time 2 was higher compared to Oravainen and Punkaharju and a similar, although statistically non-significant, trend was observed also in the *CONSTANS-like 2* (BPEV02_C10G01710) homolog (Fig. S7). In *Populus*, *APRR1/TOC1* sets the critical day length together with the *LHY* homologs and therefore regulates the timing of growth cessation (Ibáñez *et al*., 2010)*. CONSTANS*-like genes have also been associated with regulation of growth and flowering time in different forest tree species (Böhlenius *et al*., 2006; Fabbrini *et al*., 2012; Alberto *et al*., 2013b) and have been reported to be under selection in different red oak species (Lind-Riehl *et al*., 2014). In addition to the interface of circadian clock and downstream pathways, *CONSTANS* and *CONSTANS-like* genes function as hub genes between various environmental and developmental inputs and outputs and thus may be especially prone to be affected by selection for local adaptation in trees (Romero *et al*., 2024). Further, multiple genes of both *CONSTANS-like* (*COL*, 8 in *B. pendula* genome) and *PRR* families (5 in *B. pendula* genome) potentially allow subfunctionalization and fine-tuning of response in a clinal situation where differences between close-by populations’ optima are small (Lagercrantz & Axelsson, 2000; Niu *et al*., 2022; Zang *et al*., 2026).

Note, however, that due to experimental design limits, as each sampling time was represented by a single time point shortly after dawn, our ability to assess true circadian regulation, e.g., potential phase shifts between populations and expression dynamics throughout the day, was limited. Consequently, genes with expression peaks at other times of the day, including *FT*, are likely underrepresented in our analyses.

### Gene expression and co-expression indicated low pleiotropy of adaptive differences

Co-expression analysis revealed an interesting pattern: even though module eigengene expression depended largely on photoperiod and treatment duration for most modules, genes with population level differences in expression were less likely to be co-expressed with other genes indicative of underlying genetic variants having low pleiotropy.

Our results are in line with the cost of complexity theory predicting that adaptive changes close to the fitness optimum proceed through small effects with little pleiotropic effects (Orr, 2000; Wagner & Zhang, 2011). This is in contrast to mutations with high pleiotropy that may be favoured in cases where the populations are far from the optimum, as seen in a case of locally adapted common ragweed populations (Hämälä *et al*., 2020), alternative life history forms in the silverleaf sunflower (Okinedo & Moyers, 2026), or in flowering time mutations of *Arabidopsis thaliana* (Taylor *et al*., 2019), for example.

Population genetic theory predicts that as populations approach their fitness optimum, the mutations favoured by selection tend to have smaller and less pleiotropic effects, because their modest fitness benefits must outweigh the fitness costs associated with pleiotropy (Orr, 1998, 2000; Wang *et al*., 2010; Connallon & Hodgins, 2021), leading to a highly polygenic architecture of adaptive traits. Our findings therefore align with the genetic architecture of typical adaptive traits of forest trees, which are known to be under polygenic control, and previous population genomic scans that have not identified large effect adaptive loci despite strong signal of local adaptation in adaptive traits (Alberto *et al*., 2013a).

Based on a dense clinal sampling and exome-capture re-sequencing of *B. pendula*, Herrera-Egoavil *et al*., (2026) identified a putative inversion in chromosome 1 enriched for signatures of local adaptation. Curiously, the same genomic region did not stand out from the genomic background in terms of population specific gene expression in our data. There are multiple possible reasons for this discrepancy: 1) Genomic scans reveal impact of selection across the tree’s whole life cycle and tissues, across multiple generations. Our sampling is focused on single tissue in controlled conditions and limited time points and therefore does not capture all possible adaptive differences in gene expression. 2) Adaptive variation effect on gene expression is not bound to the corresponding genomic region. Even though the adaptive variation on the DNA level is at a given genomic location its impacts can be more diluted across the genome through *trans* effects of e.g., transcription factors, and 3) inversion polymorphism creates strong linkage disequilibrium of correlated allele frequency changes of which most could be essentially neutral not affecting any trait or gene expression level.

Also, Salojärvi *et al*., (2017) identified orthologs of six *Arabidopsis thaliana* genes showing clinal variation with environmental variables in *B. pendula* but none of these genes were differentially expressed between populations in our experiment. However, the analysis of Salojärvi *et al*., (2017) was conducted conditional on range-wide sweep signal, a test that is not specifically designed for identification of signals of local adaptation.

It is noteworthy that polygenic adaptation is hard to identify at the genomic level and a large proportion of it may be missed by comparative analysis relying on analysis identifying individual adaptive loci (Whiting *et al*., 2024). In the absence of pleiotropic effects, it is likely that adaptive genetic changes underlying gene expression differences are enriched in the same genes or their *cis*-regulatory areas (Stern & Orgogozo, 2008; Vande Zande & Wittkopp, 2022). Thus, the loci identified here can serve as candidates or as a proxy for genomic regions underlying polygenic adaptation. In the future, systems biology approaches connecting analysis of genome-wide diversity, transcriptomes and network level analysis should be used to obtain holistic understanding of the genetic basis of local adaptation especially when populations’ optima are close to each other.

## Supporting information

Figures S1-S7, table legends for Tables S1-S11

## Acknowledgements

We thank Soile Alatalo, Chedly Kastally, Robert Kesälahti, Alina Niskanen, Hannele Parkkinen and Olli Sarala for their help in collecting samples. We are grateful to Jarkko Salojärvi and James Ord for providing early access to the *B. pendula* and *B. pubescens* genomes. We wish to acknowledge CSC – IT Center for Science, Finland, for computational resources.

This work was funded by the Research Council of Finland (grant numbers 346139, 329438 and 329440).

## Author contributions

TP, JV, KK and SK planned the study. JV conducted the controlled growth chamber experiments. JP, YP and JS conducted the RNA extractions. JS and YP performed the computational analysis. KK and SK provided the material. JS and TP wrote the manuscript. TP and JPV supervised analysis and writing. All authors read and approved the final manuscript.

## Data availability

All sequencing data have been deposited in the NCBI Sequence Read Archive (SRA) under the accession code PRJNA1537397. The processed gene count matrix, seedling growth data, and R scripts used for data processing, statistical analyses, and figure generation are available on Figshare at https://doi.org/10.6084/m9.figshare.c.8756214. All accession numbers and repository links will be made publicly available upon publication.

