## Supplementary material for "Gene expression dynamics through time reveal low pleiotropy genes underlying local adaptation in silver birch (*Betula pendula*)": Figures S1-S7, table legends for Tables S1-S11

**Fig. S1** (a) Sample clustering before outlier removal. (b) Principal component analysis (PCA) of variance-stabilized gene expression data from Betula pendula seedlings before outlier removal. Each point represents a sample, positioned according to the first two principal components (PC1, PC2).

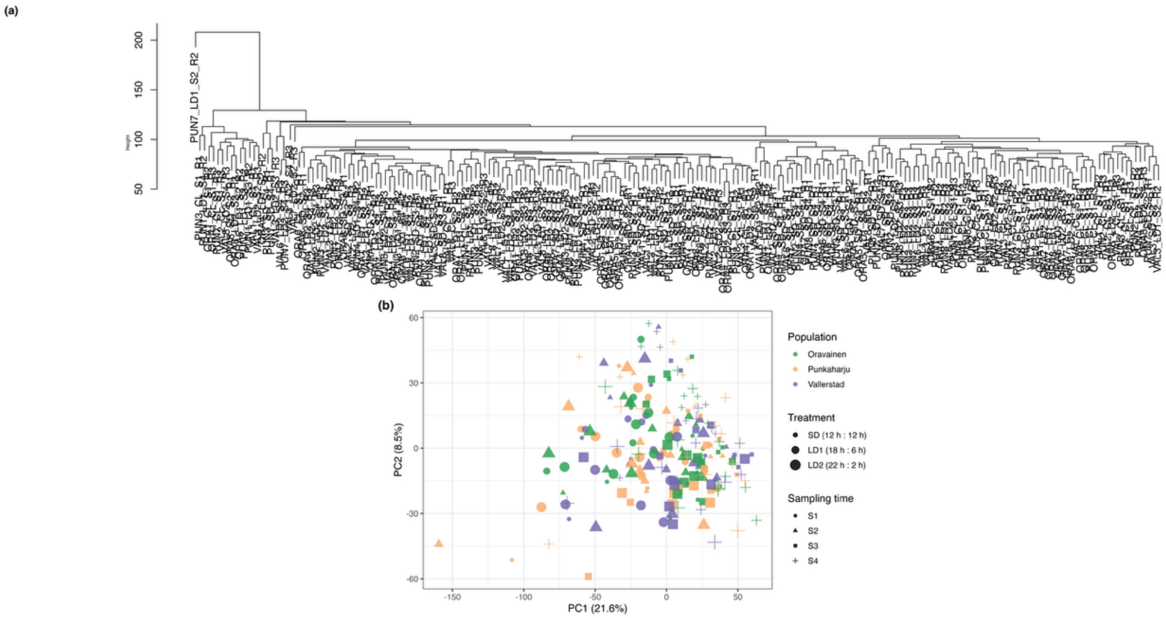


**Fig. S2** (a) Observed absolute growth rates of Betula pendula seedlings. Boxplots represent samples from different populations. Panels represent treatments. (b) Predicted growth rate for each population under different photoperiod treatments. Points represent absolute growth rates for each sample. Lines represent estimated marginal means derived from the fitted interaction model, and shaded ribbons indicate 95% confidence intervals. Panels are faceted by photoperiod treatment, colours denote population, x-axis represents sampling time (days since sowing) and y-axis shows the predicted growth rates for each population.

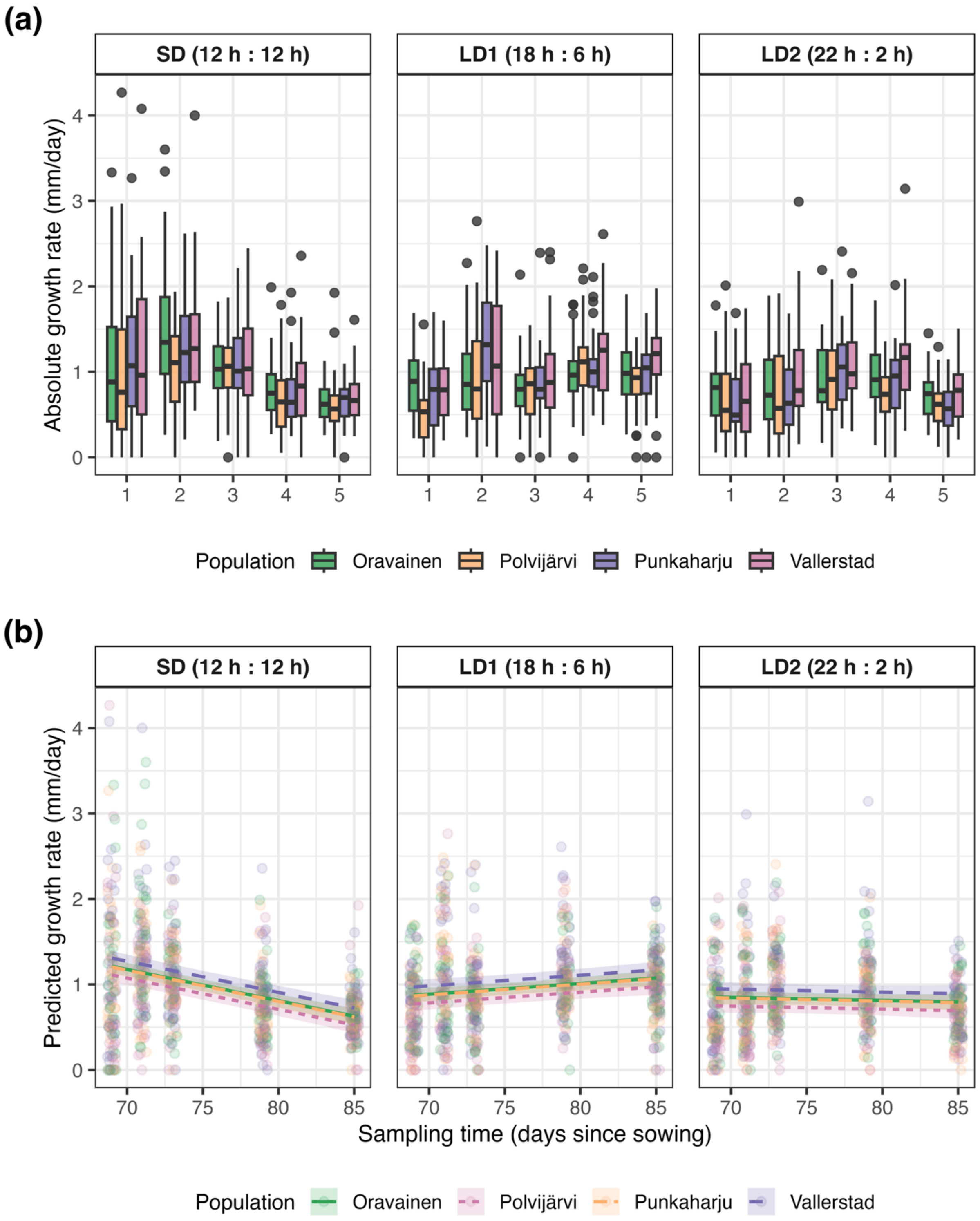
(a)
SD (12 h : 12 h)
LD1 (18 h : 6 h)
LD2 (22 h : 2 h)
Absolute growth rate (mm/day)
0
1
2
3
4
1
2
3
4
5
Population
Oravainen
Polvijärvi
Punkaharju
Vallerstad
(b)
SD (12 h : 12 h)
LD1 (18 h : 6 h)
LD2 (22 h : 2 h)
Predicted growth rate (mm/day)
70
75
80
85
Sampling time (days since sowing)
Population
Oravainen
Polvijärvi
Punkaharju
Vallerstad

**Fig. S3** Effect sizes of the growth model for Betula pendula seedlings. Point estimates and 95% confidence intervals for the fixed effects from the linear mixed-effects model fitted to estimate absolute growth rates. Population Oravainen in SD treatment was used as the baseline.

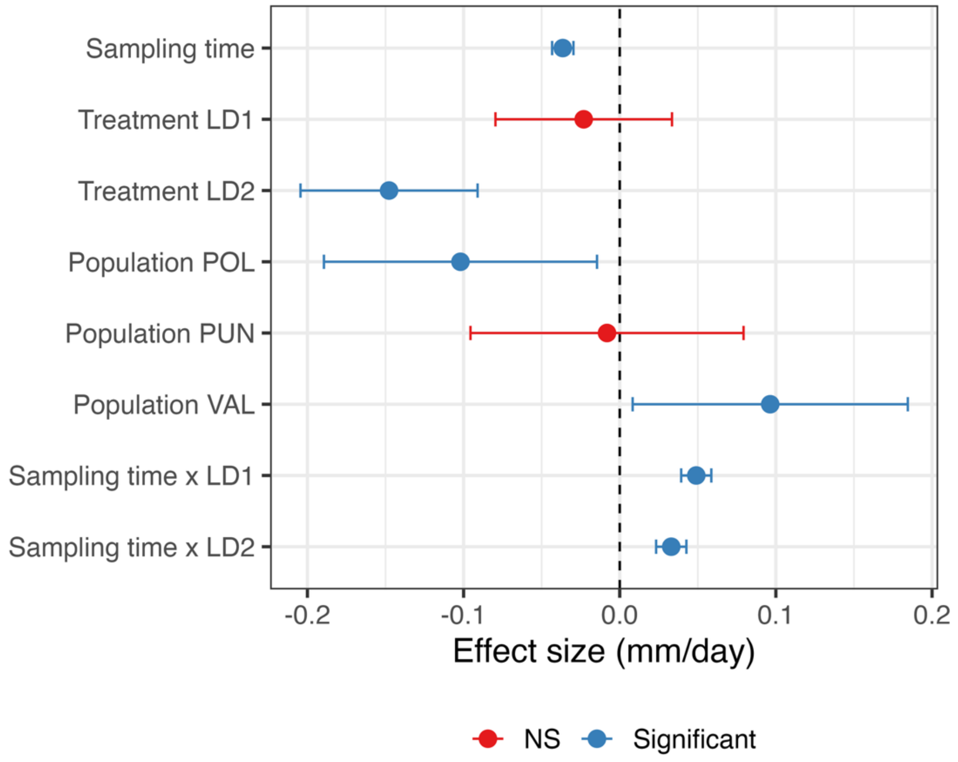


**Fig. S4** Multimapping to Betula pubescens (polyploid) vs Betula pendula (diploid). Each point represents a sample and colours represent populations.

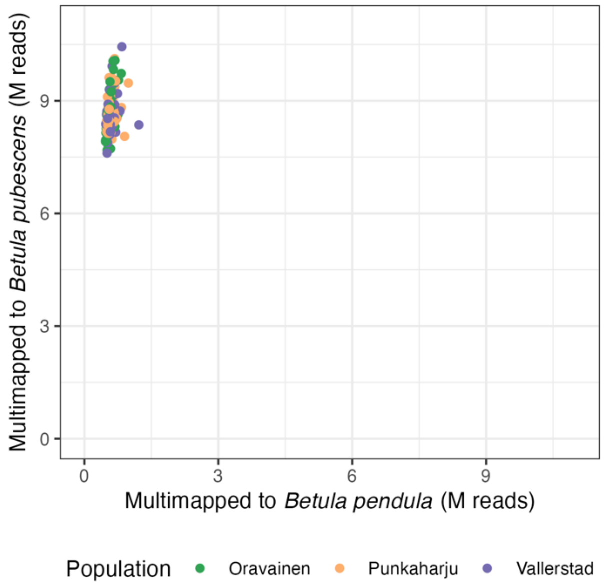


**Fig. S5** Relationship of the co-expression module eigengene expression to the treatments in Betula pendula seedlings. Heatmap showing regression coefficients from linear models fitted to module eigengenes (MEs) of weighted gene co-expression network analysis (WGCNA) modules. Each panel has a different population at sampling time 1 in constant light (SD room) as a baseline. Each cell of the heatmap represents the estimated effect of a model term (x-axis) on a given module eigengene (y-axis). Colour indicates the direction and magnitude of the effect (blue, negative; white, no effect; red, positive). Asterisks denote false discovery rate adjusted significance levels (***: p ≤ 0.001, **: p ≤ 0.01, *: p ≤ 0.05).

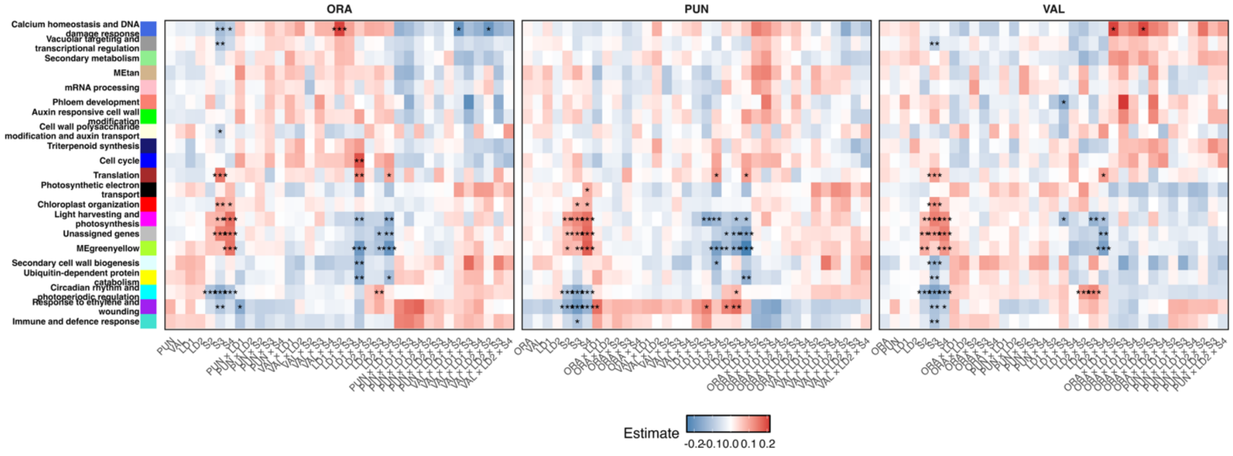


**Fig. S6** Module eigengene expression across populations and treatments. Weighted co-expression network analysis (WGCNA) module eigengene (ME) expression across time points.

Lines indicate the mean eigengene value for each population across sampling time points. Panels are faceted by treatment. Each subplot corresponds to a single co-expression module with the background colour corresponding to the colour of the module and the header to the name derived from Gene Ontology enrichment analysis.

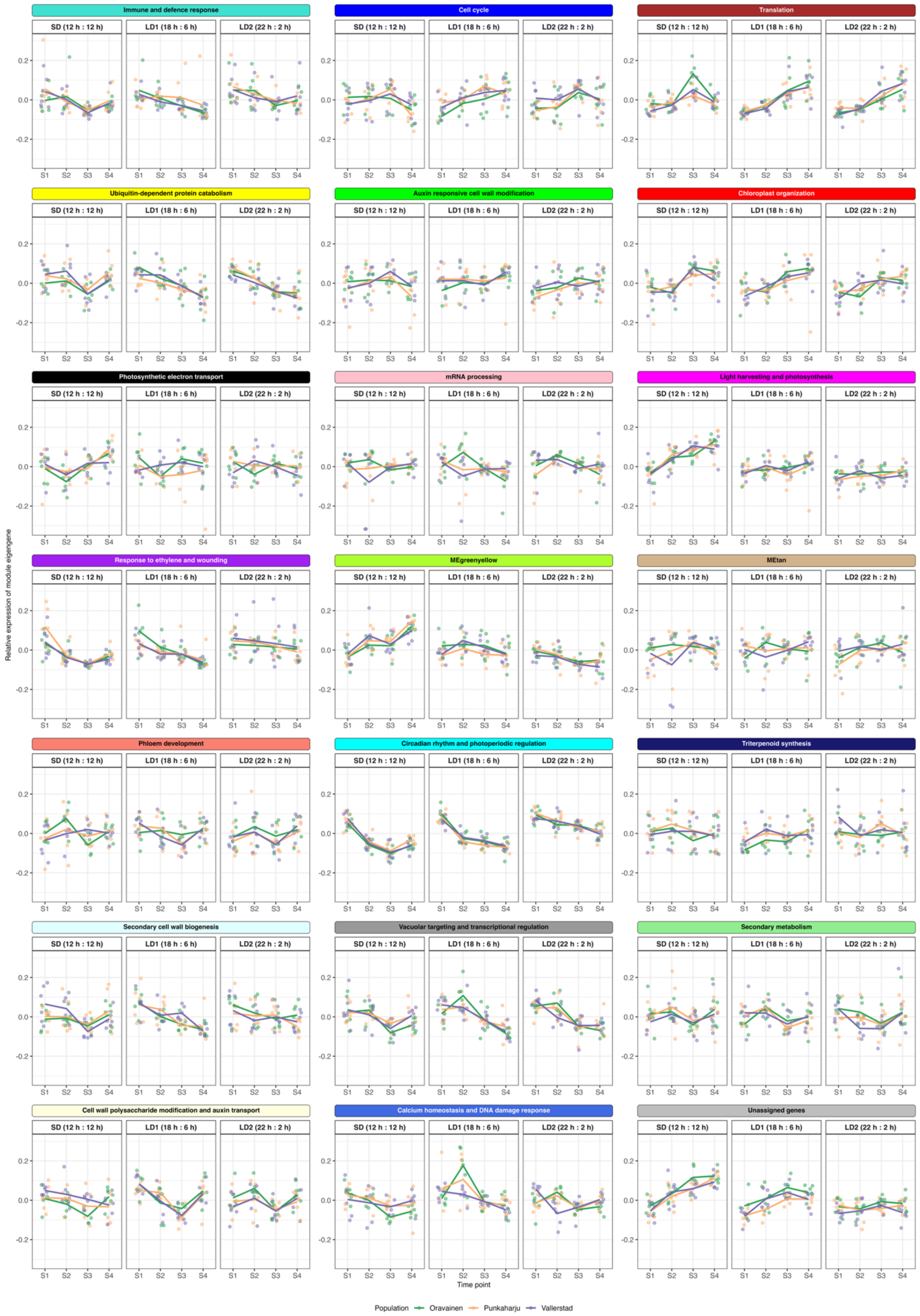
Immune and defence response
Cell cycle
Translation
Ubiquitin-dependent protein catabolism
Auxin responsive cell wall modification
Chloroplast organization
Photosynthetic electron transport
mRNA processing
Light harvesting and photosynthesis
Response to ethylene and wounding
MEgreenyellow
MEtan
Phloem development
Circadian rhythm and photoperiodic regulation
Triterpenoid synthesis
Secondary cell wall biogenesis
Vacuolar targeting and transcriptional regulation
Secondary metabolism
Cell wall polysaccharide modification and auxin transport
Calcium homeostasis and DNA damage response
Unassigned genes
SD (12 h : 12 h)
LD1 (18 h : 6 h)
LD2 (22 h : 2 h)
0.2
0.0
-0.2
S1
S2
S3
S4
Relative expression of module eigengene
Time point
Population
Oravainen
Punkaharju
Vallerstad

**Fig. S7** Circadian clock and flowering time gene expression in Betula pendula seedlings. Lines indicate population averages in expression with vertical bars indicating standard error in expression within populations. The x-axis differs between plots.

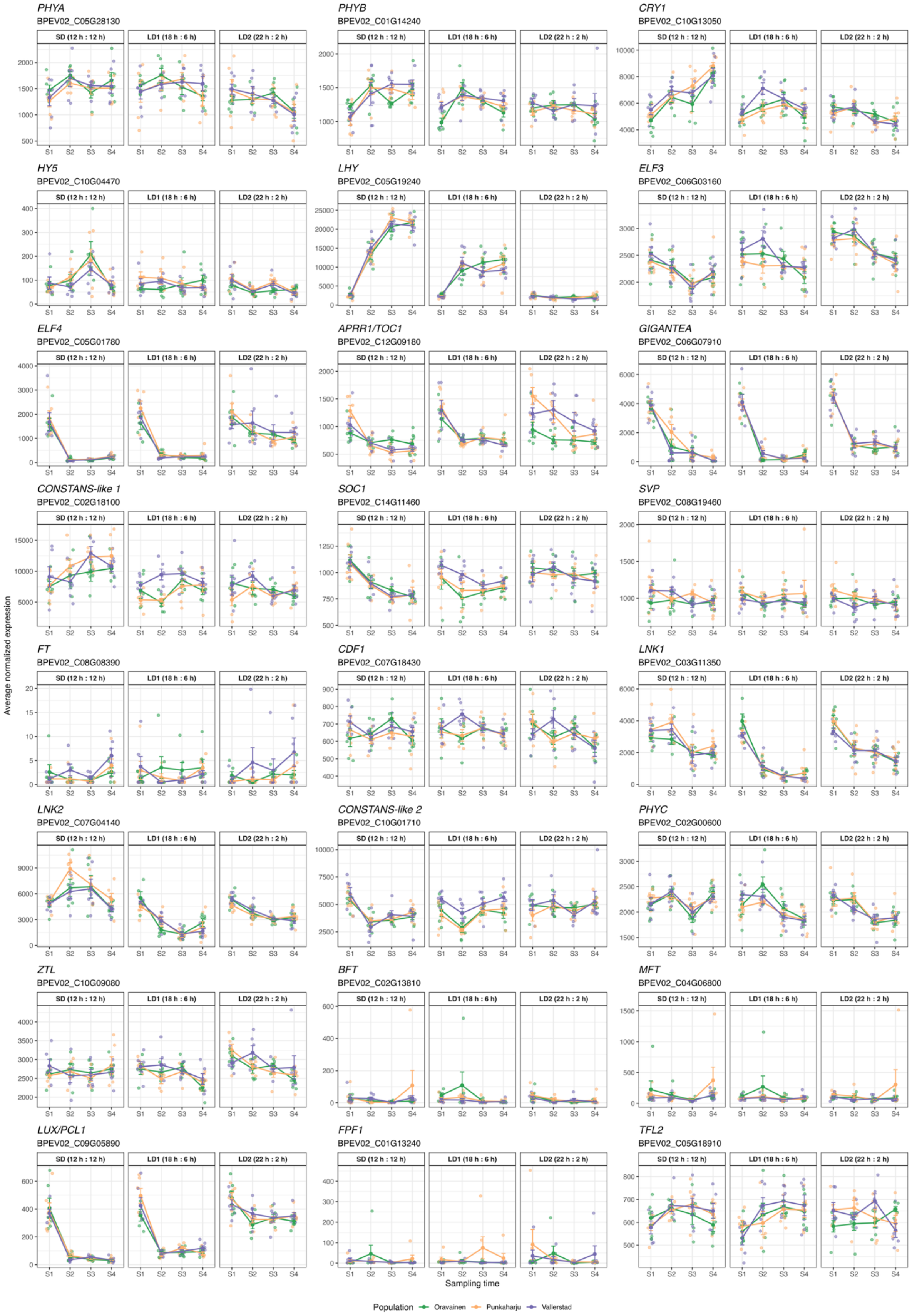
PHYA
BPEV02_C05G28130
PHYB
BPEV02_C01G14240
CRY1
BPEV02_C10G13050
HY5
BPEV02_C10G04470
LHY
BPEV02_C05G19240
ELF3
BPEV02_C06G03160
ELF4
BPEV02_C05G01780
APRR1/TOC1
BPEV02_C12G09180
GIGANTEA
BPEV02_C06G07910
CONSTANS-like 1
BPEV02_C02G18100
SOC1
BPEV02_C14G11460
SVP
BPEV02_C08G19460
FT
BPEV02_C08G08390
CDF1
BPEV02_C07G18430
LNK1
BPEV02_C03G11350
LNK2
BPEV02_C07G04140
CONSTANS-like 2
BPEV02_C10G01710
PHYC
BPEV02_C02G00600
ZTL
BPEV02_C10G09080
BFT
BPEV02_C02G13810
MFT
BPEV02_C04G06800
LUX/PCL1
BPEV02_C09G05890
FPF1
BPEV02_C01G13240
TFL2
BPEV02_C05G18910
SD (12 h : 12 h)
LD1 (18 h : 6 h)
LD2 (22 h : 2 h)
S1
S2
S3
S4
Average normalized expression
Sampling time
Population
Oravainen
Punkaharju
Vallerstad

**Table S1** Sample summary table.

**Table S2** Circadian clock and flowering time gene homologs used in this study.

**Table S3** Growth model comparison with AIC and BIC.

**Table S4** Top genes for which the modelled factors explain the majority of expression variance.

**Table S5** List of genes assigned to each co-expression module.

**Table S6** Gene Ontology enrichment results for each co-expression module.

**Table S7** Variance partition for co-expression modules.

**Table S8** Gene lists from likelihood ratio tests annotated with co-expression module metrics.

**Table S9** Continuation for Table 4, Fisher's exact pairwise test. Odds ratio (OR) < 1: genes are more likely to be assigned to a co-expression module; OR >1: genes are more likely to be unassigned to any co-expression module.

**Table S10** Summary of differential gene expression results in relation to weighted co-expression network analysis (WGCNA). Average connectivity (kME) is calculated over all modules.

**Table S11** Circadian clock and flowering time gene weighted co-expression network module membership and connectivity. kME measures how strongly a gene correlates with the module eigengene. kWithin measure how strongly a gene is connected to other genes in the same module. Hub percentile scales the kWithin to the module size.
